# Structural and compositional profiling of individual biomolecular assemblies by infrared absorbance-modulated evanescent scattering

**DOI:** 10.64898/2026.03.12.711458

**Authors:** Qing Xia, Qingbo Wang, Danchen Jia, Dashan Dong, Mingsheng Li, Eliana Sherman, Jianpeng Ao, Qian Ren, Fiamma Ayelen Buratti, Pernilla Wittung-Stafshede, Rylie Bolarinho, Huan Bao, Lulu Jiang, Ji-Xin Cheng

**Affiliations:** Department of Electrical and Computer Engineering, Boston University, Boston, Massachusetts 02215, United States; Photonics Center, Boston University, Boston, Massachusetts 02215, United States; Department of Neuroscience, University of Virginia School of Medicine, Charlottesville, Virginia 22908, United States; Center for Brain Immunology and Glia (BIG), University of Virginia School of Medicine, Charlottesville, Virginia 22908, United States; Department of Molecular Physiology and Biological Physics, University of Virginia, Charlottesville, Virginia 22903, United States; Department of Chemistry, Rice University, Houston, Texas 77005, United States; Department of Chemistry, Boston University, Boston, Massachusetts 02215, United States; Department of Biomedical Engineering, Boston University, Boston, Massachusetts 02215, United States

## Abstract

The structure and composition of biomolecular assemblies shape their biological functions and disease processes. Yet existing approaches either require homogeneous samples or lack chemical specificity and sensitivity, limiting the elucidation of transient oligomeric species under native conditions. Here we report infrared absorbance-modulated evanescent scattering (IR-AMES) that enables ultrasensitive structural and compositional profiling of individual biomolecular assemblies in aqueous environments. This capability is achieved by photothermally encoding mid-infrared vibrational absorption into scattering contrast using a total-internal-reflection geometry. Applying IR-AMES to recombinant tau assemblies, we resolve random-coil monomers and the emergence of heterogeneous oligomers beyond ensemble-averaged measurements. In Alzheimer’s disease brain–derived tau, IR-AMES reveals enrichment of antiparallel β-sheet structures and RNA components in oligomers. Using lipid nanodiscs as membrane mimics, we find that pathological tau oligomers exhibit enhanced interactions with anionic membranes. IR-AMES enables spectroscopic imaging of diverse biomolecular assemblies, opening opportunities to elucidate structure–function relationships in heterogeneous biomolecular systems.

---

Biomolecular assemblies, particularly intrinsically disordered proteins, exist as structurally and chemically heterogeneous ensembles that underlie their diverse biological functions and disease processes^1, 2^. Resolving the structure and composition at the single-assembly level under native conditions remains challenging, especially for transient oligomeric species that are obscured by ensemble-averaged measurements^3^. Established structural biology techniques, including X-ray crystallography^4^, cryo-electron microscopy^5–7^ and AI-based structure prediction^8, 9^, rely on homogeneous or static conformations, thus limiting the structural characterization of heterogeneous protein assemblies.

Optical and scanning-probe approaches have enabled the detection of individual biomolecules, yet each provides only a partial view of heterogeneous assemblies. Methods that probe single-molecule conformations, including high-speed atomic force microscopy^10^ and fluorescence-based techniques^11, 12^, reveal dynamic heterogeneity but lack intrinsic chemical or structural information. Surface-enhanced vibrational spectroscopies provide chemical contrast yet suffer from hotspot-dependent variability^13–15^. Near-field infrared (IR) nanospectroscopy resolves protein secondary structures with nanoscale resolution, but is typically limited to large fibrils or dried oligomers and has low throughput^16–18^. Far- field interferometric scattering enables high-sensitivity, high-throughput detection of single proteins in solution, yet lacks chemical contrast^19–22^. As a result, resolving the structure and composition of individual oligomeric assemblies under native conditions remains a major challenge.

Here we introduce infrared absorbance-modulated evanescent scattering (IR-AMES) that enables ultrasensitive chemical fingerprinting of individual biomolecular assemblies in aqueous environments. IR-AMES photothermally encodes mid-infrared vibrational absorption into scattering contrast in a total-internal-reflection geometry, allowing chemically specific and structure-sensitive characterization of individual assemblies. We first benchmark IR-AMES using polymer nanoparticles to quantify detection sensitivity and spatial resolution. We then apply IR-AMES to protein assemblies in buffer, using immunoglobulin M (IgM) as a model oligomeric system to resolve hydration-dependent conformational heterogeneity in individual protein complexes. Extending this approach to the intrinsically disordered protein tau, IR-AMES resolves the structure and composition of tau assemblies across aggregation states and associated functional properties such as membrane interaction and cytotoxicity. Beyond tau, we further demonstrate the generality of IR-AMES by characterizing structurally defined α-synuclein amyloid fibrils, highlighting its applicability to diverse biomolecular assemblies. Together, these results establish IR-AMES as a nanoscale spectroscopic imaging platform for linking molecular structure and composition to biological function in complex biomolecular assemblies.

## IR-AMES principle, setup and benchmarks

IR-AMES operates by converting mid-infrared vibrational absorption into a modulated evanescent scattering signal via the photothermal effect^23, 24^. As illustrated in **Fig. 1a**, IR excitation of molecular vibrations (e.g., amide C=O stretching) generates local heating that produces a transient temperature increase (Δ*T*), which leads to thermal expansion (Δ*r*), and refractive index decrease (Δ*n*) of the molecule. These perturbations are detected in a pump-probe scheme as a modulation of the evanescent scattering intensity (Δ*I*), which depends on the local probe- and pump-field intensities, while the detection sensitivity is determined by the modulation depth (Δ*I*/*I*).

**Fig. 1.**
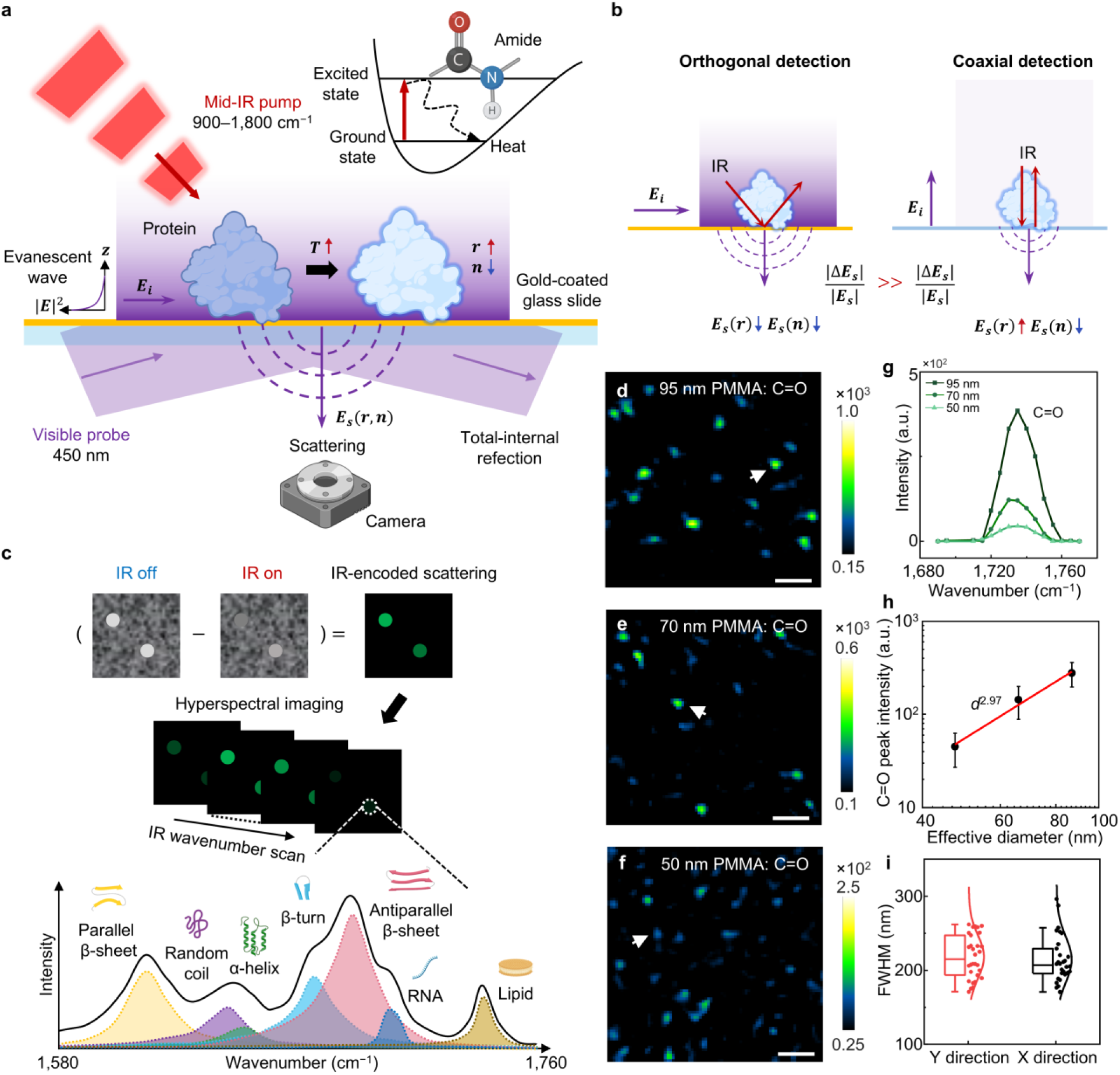
IR-AMES concept, implementation and performance. a, Schematic of IR-AMES. Mid-IR excitation of molecular vibrations (e.g., amide modes in proteins) induces non-radiative relaxation, generating localized heating that transiently modulates the particle radius (*r*) and refractive index (*n*), thereby perturbing the scattered field from the sample (*E_s_*). A visible probe undergoes total internal reflection (TIR) at a gold-coated glass substrate to generate a surface-confined evanescent incident field (*E_i_*). A pulsed mid-IR pump is introduced at an oblique angle to excite molecular vibrations. Evanescent scattering is collected orthogonally for wide-field detection. *T*: temperature. **b,** Conventional photothermal detection uses a coaxial geometry (right), where photothermal induced scattering field cancellation limits modulation depth 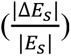. In contrast, IR-AMES employs an orthogonal detection scheme of TIR-illuminated evanescent scattering (left), which enhances the relative scattering modulation. **c,** Image-processing workflow. IR-encoded scattering images are generated by subtracting IR-off from IR-on frames. IR wavenumber scanning produces hyperspectral stacks for single-particle compositional and structural analysis. **d–f,** IR-AMES images of 95-nm (**d**), 70-nm (**e**) and 50-nm (**f**) PMMA particles at the C=O band (1,735 cm^−1^). Scale bars: 1 µm. **g,** Representative single-particle C=O stretching spectra corresponding to particles marked by white arrows in **d–f**. **h,** IR-AMES C=O signal intensity versus effective particle diameter (*d*) under the evanescent field. Data points represent mean values, and error bars indicate the full width at half maximum (FWHM) of Gaussian fits to the corresponding intensity distributions. **i**, Spatial resolution of IR-AMES, quantified from the FWHM of 50-nm PMMA nanoparticles in integrated C=O-band images. Mean FWHM values are 218 nm and 214 nm along the *y* and *x* directions (*n* = 30 particles), respectively.

Earlier efforts^25, 26^ using high-energy optical parametric oscillator (OPO) mid-IR sources and interferometric enhancement pushed mid-IR photothermal sensitivity to single ∼100-nm viruses in air. However, the conventional coaxial photothermal detection (**Fig. 1b**, right) faces an intrinsic physical limit to its sensitivity: Δ*r* and Δ*n* have opposite contributions to scattering intensity modulation^26^, reducing the overall modulation depth. Thus, breaking the single-molecule barrier requires a more efficient photothermal detection strategy.

To overcome this limitation, IR-AMES boosts the sensitivity by an orthogonal detection scheme with evanescent-field–enhanced readout (**Fig. 1b**, left, details in **Supplementary Note 1**). First, an IR- reflective gold-coated substrate with 45° *p*-polarized mid-IR incidence doubles the interfacial pump field intensity (**Supplementary Fig. 1**). Second, a surface-confined evanescent probe field enhances the interferometric scattering by ∼6.5-fold (**Supplementary Fig. 2**). Third, the horizontally propagating evanescent wave allows the scattering to be collected orthogonally (**Supplementary Fig. 3**). Together, this scheme eliminates cancellation between the two photothermal contributions (**Fig. 1b**, left) and boosts the modulation depth over coaxial detection (**Fig. 1b**, right).

The experimental implementation is illustrated in **Fig. 1a** (Details in **Supplementary Fig. 4** and **Methods**). A visible probe beam undergoes total internal reflection (TIR) on a gold-coated substrate to generate a surface-confined evanescent field, which enhances scattering from interfacial molecules and enables orthogonal collection of the scattered signal. To encode vibrational spectra, a pulsed mid-IR pump beam illuminates the sample at an oblique angle. The reflective gold film doubles the local IR excitation while minimizing unwanted heating of the immersion oil. The mid-IR pump, visible probe, and camera are electronically synchronized (**Supplementary Fig. 5**) to ensure consistent acquisitions of IR-on and IR-off images. The IR absorption induces transient photothermal modulation of the scattering signal, and subtraction of IR-on and IR-off frames produces an IR-encoded scattering image. By scanning the IR wavenumber, a hyperspectral image stack is generated, from which vibrational fingerprints containing molecular composition and structure signatures are obtained (**Fig. 1c** and **Supplementary Fig. 6**).

We characterized the photothermal signal enhancement in the orthogonal detection geometry using 500-nm poly(methyl methacrylate) (PMMA) beads, a standard benchmark used in coaxial scattering-based photothermal microscopy owing to their strong C=O vibrational resonance. These particles show strong resonant contrast at 1,728 cm^−1^, with negligible off-resonance signal (**Extended Data Fig. 1a**). The photothermal modulation depth reaches ∼22.7% for a single bead in IR-AMES (**Extended Data Fig. 1b**), whereas coaxial photothermal detection produces only ∼0.2%^27^. This >100-fold sensitivity enhancement enables single-shot chemical-bond imaging at 100 Hz, with the potential extension to kilohertz frame rates using a high-speed camera^28^. The modulation depth increases linearly with IR power (**Supplementary Fig. 7**), confirming minimal nonlinear thermal artifacts. Despite operating with a quantum-cascade laser (QCL) delivering 10-fold lower pulse energy than an IR OPO and a camera 6- fold slower in frame rate, IR-AMES achieves ∼60-fold faster chemical-bond imaging than earlier OPO- based interferometric mid-IR photothermal implementations^25^ (**Supplementary Table 1**). A broader comparison with representative optical photothermal infrared (O-PTIR) implementations is provided in **Supplementary Note 5** and **Supplementary Fig. 8**. A hyperspectral image stack spanning 900– 1,800 cm^−1^ with a 2 cm^−1^ step size (450 spectral points) is acquired within two minutes, primarily limited by the QCL tuning speed. The statistical spectra of 50 individual PMMA beads match Fourier-transform infrared spectroscopy (FTIR) results well and clearly resolve the C–O–C and C=O vibrational modes, with a small standard deviation of 3.05×10^-2^ across all wavenumbers (**Extended Data Fig. 1c**). To further assess spectral fidelity at smaller particle sizes, we performed correlative total internal reflection fluorescence (TIRF) and IR-AMES measurements of fluorescent 38-nm polystyrene (PS) nanoparticles. The two modalities showed consistent particle distributions, and the single-particle IR-AMES spectra exhibited the characteristic C–H and aromatic C=C bands of PS, consistent with bulk FTIR measurements (**Extended Data Figs. 1d–f**). Atomic force microscopy (AFM) measurements further confirmed the morphology and size distributions of the standard nanoparticles used in these experiments (**Supplementary Fig. 9**).

Size-dependent photothermal responses were quantified using PMMA beads of 95, 70, and 50 nm diameter (**Figs. 1d–f**). The C=O peak intensity decreases with particle size, and the mean image intensity scales with the third power of diameter (**Fig. 1g, h** and **Supplementary Fig. 10**), consistent with the interferometric scattering theory^29^ after accounting for the evanescent-field decay as an effective diameter (**Supplementary Note 2**). The interferometric reference field arises from nanoscale roughness of the gold film, consistent with mechanisms established previously^21^. The lateral spatial resolution was characterized using 50-nm particles. Analysis of 30 particles yielded mean full width at half maximum (FWHM) values of 218 nm and 214 nm along the *y* and *x* directions, respectively (**Fig. 1i**), with representative line-profiles shown in **Supplementary Fig. 11**. IR illumination uniformity was validated by beam-profile fitting, and a central 30-µm region was selected as the effective imaging area (**Supplementary Fig. 12**). Under the applied excitation conditions, thermal modelling and experimental validation estimate a transient temperature rise of ∼36.7 K for 500-nm PMMA particles. For 50-nm PMMA particles, the temperature quickly rises to ∼0.41 K on the nanosecond scale and remains constant during IR excitation (**Supplementary Notes 3–4**, **Supplementary Figs. 13–14**). Such small temperature rise is compatible with biomolecular measurements. Together, these results establish IR-AMES as a label-free, high-throughput and high-fidelity platform for quantitative vibrational spectroscopy of individual nano-objects.

## Vibrational fingerprinting of individual protein complexes in buffer

To establish vibrational fingerprinting of individual protein complexes, we applied IR-AMES to IgM (∼950 kDa), an oligomeric protein that comprises both β-sheet and conformationally flexible regions^30^. As an initial benchmark, IgM was measured after drying on the substrate. AFM confirmed the presence of isolated IgM complexes^31^ with lateral dimensions of ∼35 nm and heights of ∼5 nm (**Supplementary Fig. 15**). IR-AMES imaging resolves discrete diffraction-limited spots (**Fig. 2a**), and spectra extracted from these spots show characteristic protein vibrational features across the fingerprint region, including the amide-I and amide-II bands. The mean IR-AMES spectrum closely agrees with the corresponding FTIR spectrum of dried IgM (**Supplementary Fig. 16**). We then focused on the amide-I band to quantify relative protein content and probe structural signatures through its secondary-structure- sensitive backbone C=O stretching vibration^32^. The amide-I spectra exhibit three quantized intensity levels (**Fig. 2b**), corresponding to one, two, and three IgM complexes within the detection area. Histogram analysis of ∼300 particles reveals a multimodal intensity distribution (**Fig. 2c**). Gaussian fitting shows that the peak centers scale linearly with molecular count (**Fig. 2d**), supporting detection at the single-complex level^20^.

**Fig. 2.**
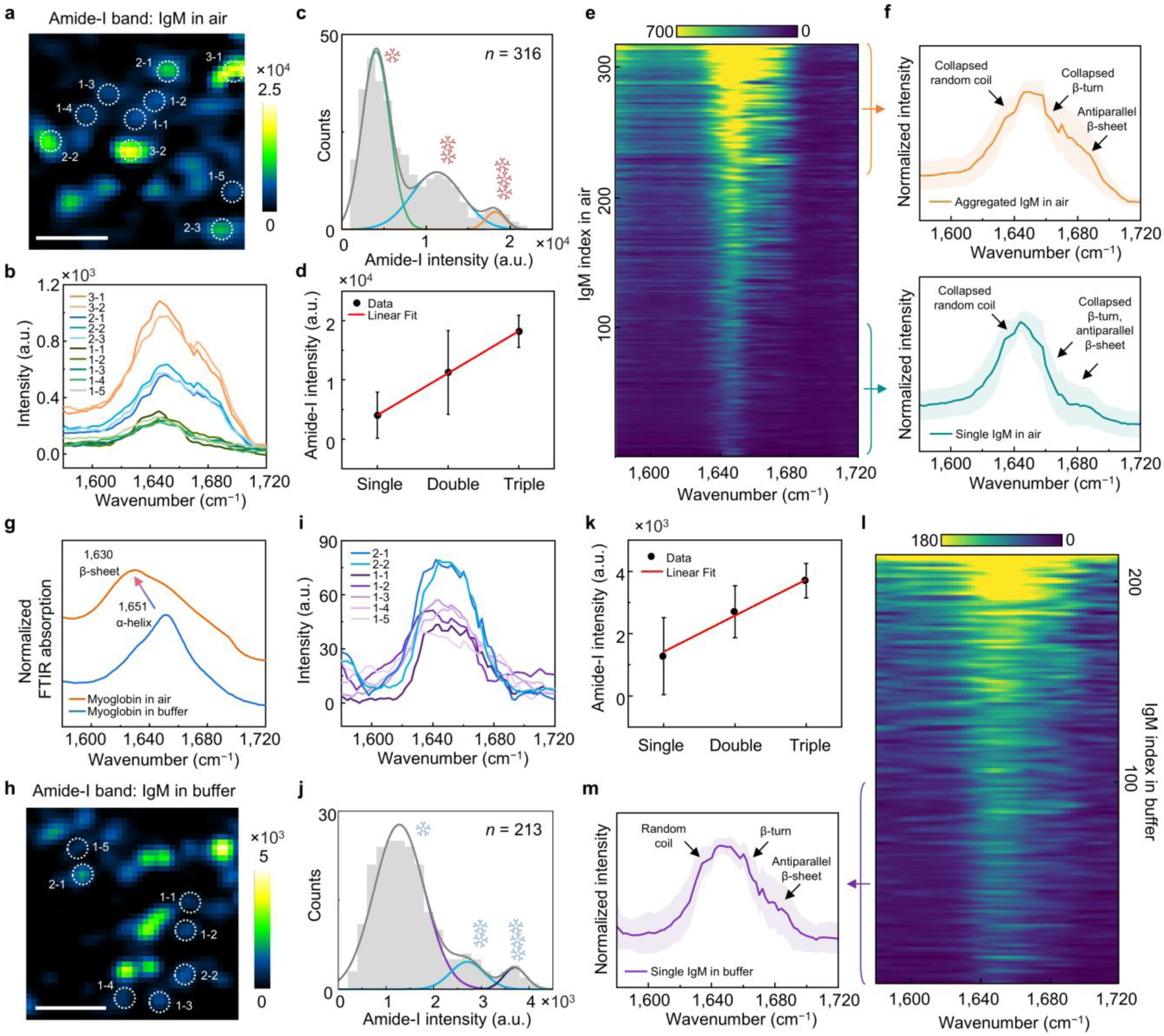
Fingerprinting single protein complexes in air and buffer. a–f, IgM in air. **a,** IR-AMES image of IgM molecules at the amide-I band. Scale bar: 1 µm. **b,** Representative IR-AMES spectra extracted from circled IgM molecules in **a**, showing increased amide-I intensity. **c,** Amide-I intensity histograms of IgM molecules with Gaussian fits. Secondary (blue) and tertiary (orange) peaks correspond to multi-molecule binding or multiple molecules within sub-diffraction distances. **d,** IR-AMES intensity versus IgM molecule count. Data points represent the mean of each peak in **c**. Error bars indicate the fitted Gaussian FWHM. **e,** Heatmap of IR-AMES spectra in the amide-I region, sorted by integrated intensity over amide-I band. **f,** Representative average spectra for aggregated IgM (index 217–316) and single IgM (index 1–100) in air. Solid line: mean spectra, shaded region: standard deviation. **g,** Dehydration-induced distorts of native myoglobin secondary structures, with FTIR spectra shifting from α-helix (buffer, 20 mg/ml) to β-sheet (air, powder). Spectra were normalized to 0–1 and vertically offset for clarity. **h–m,** IgM in buffer. **h,** IR-AMES image of IgM molecules at the amide-I band. Scale bar: 1 µm. **i,** Representative IR-AMES spectra extracted from circled IgM molecules in **h**. **j,** Amide-I intensity histograms of IgM molecules with Gaussian fits. Secondary (light blue) and tertiary (dark blue) peaks indicate multi-molecule binding events or multiple molecules within sub-diffraction distances. **k,** IR- AMES image intensity versus IgM molecule count extracted as in **d**. **l,** Heatmap of IR-AMES spectra of IgM, sorted by integrated intensity over amide-I band. **m,** Representative average spectra for single IgM molecules in buffer (index 1–100) in **l**. Solid line: mean spectra, shaded region: standard deviation.

IR-AMES spectroscopy heatmaps further reveal conformational heterogeneity among IgM molecules, with the amide-I intensity increasing with molecular count (**Fig. 2e**). Spectra from single IgM molecules exhibit partially collapsed random-coil, β-turn, and antiparallel β-sheet features induced by dehydration (**Fig. 2f**, bottom), whereas the aggregates retain stronger antiparallel β-sheet signatures due to intermolecular stabilization (**Fig. 2f**, top). We suspect that the spectral difference between single IgMs and aggregates arises from dehydration. To validate, we performed independent FTIR measurements on the more flexible, α-helix-dominated myoglobin and observed even stronger structural distortion upon dehydration, with a distinct shift toward β-sheet-rich spectra (**Fig. 2g**). These results highlight the need for single-protein IR spectroscopy under buffer conditions, where native protein conformations are preserved.

For the solution environment, however, the protein signals are easily overwhelmed by water absorption background. We addressed this challenge by using a thin aqueous layer, oblique mid-IR incidence, and short 80-ns IR pulses to suppress the water background (**Extended Data Figs. 2a–d**). Both simulations and measurements on ∼100-nm PMMA in water confirm that, under these conditions, the molecular C=O band at 1,728 cm^−1^ dominates the photothermal response. We further validated detection at smaller particle sizes by correlative TIRF and IR-AMES imaging of fluorescent 38-nm PS nanoparticles in water. The two modalities showed consistent particle distributions, while IR-AMES resolved the characteristic single-particle vibrational signatures of PS (**Extended Data Figs. 2e, f**).

With this strategy, IR-AMES detects hydrated IgM at single-complex sensitivity (**Fig. 2h, i**). The resulting intensity distribution is dominated by the lowest-intensity population, indicating that most detected events correspond to individual IgM molecules rather than small aggregates (**Fig. 2j, k**). Importantly, the hydrated IgM molecules (**Fig. 2l, m**) exhibit a broader amide-I band than the dried ones, consistent with hydration-induced stabilization of flexible random-coil, β-turn, and antiparallel β-sheet conformations^30^.

To demonstrate the ability of IR-AMES to resolve structural and compositional differences, we measured reference biomolecules with distinct vibrational signatures. IR-AMES spectra of parallel β- sheet-rich concanavalin A, α-helix-rich myoglobin and human RNA^33^ resolve characteristic protein- and RNA-associated bands in solution (**Supplementary Fig. 17** and **Supplementary Table 2).** For the reference proteins, both the mean IR-AMES spectra and second-derivative minima are consistent with the corresponding FTIR measurements, validating the assignment of characteristic secondary-structure bands (**Extended Data Fig. 3**). Together, these results establish IR-AMES as a platform capable of resolving structure and composition of individual biomolecular assemblies in solution.

## IR-AMES reveals conformational heterogeneity in recombinant tau oligomers

Having established that IR-AMES can resolve the structure and composition of individual protein assemblies in solution, we next applied this approach to more complex and heterogeneous systems. Tau, a prototypical intrinsically disordered protein associated with Alzheimer’s disease (AD)^34–36^, undergoes oligomerization from largely disordered monomers into oligomers that precede fibril formation. Increasing evidence implicates soluble tau oligomers as the primary neurotoxic species, yet the molecular features that distinguish different oligomeric states remain poorly understood^37–39^.

To resolve these structural transitions, we next performed IR-AMES measurements on recombinant human 2N4R tau monomer (rTauM) and oligomer (rTauO) samples (**Fig. 3a**). The distribution of amide- I intensities for rTauM exhibits a single Gaussian peak, indicating a largely homogeneous monomer population (**Fig. 3b**). In contrast, rTauO displays multiple discrete intensity populations. Plotting the peak centers against the apparent oligomer order reveals an approximately linear scaling (**Fig. 3c**). The lowest-intensity rTauO population lies close to the rTauM distribution, potentially reflecting residual monomeric species or small assemblies, whereas subsequent populations follow the fitted trend toward higher intensities. These distributions are consistent with rTauO assemblies that predominantly comprise dimers and trimers^40, 41^. In addition, IR-AMES spectra of rTauM are relatively homogeneous, with dominant intensity centered near 1,642 cm^−1^, and strong particle contrast is observed primarily in the random-coil channel (**Fig. 3d**). Although individual monomers are conformationally dynamic, their spectra cluster within a narrow range, indicating that random coil with minor α-helical contributions dominates at the single-particle level. This spectral uniformity is consistent with AlphaFold3 structure predictions^8^, which show highly disordered tau monomers with substantial conformational variability but no stable β-sheet formation (**Supplementary Note 6** and **Supplementary Fig. 18a**).

**Fig. 3.**
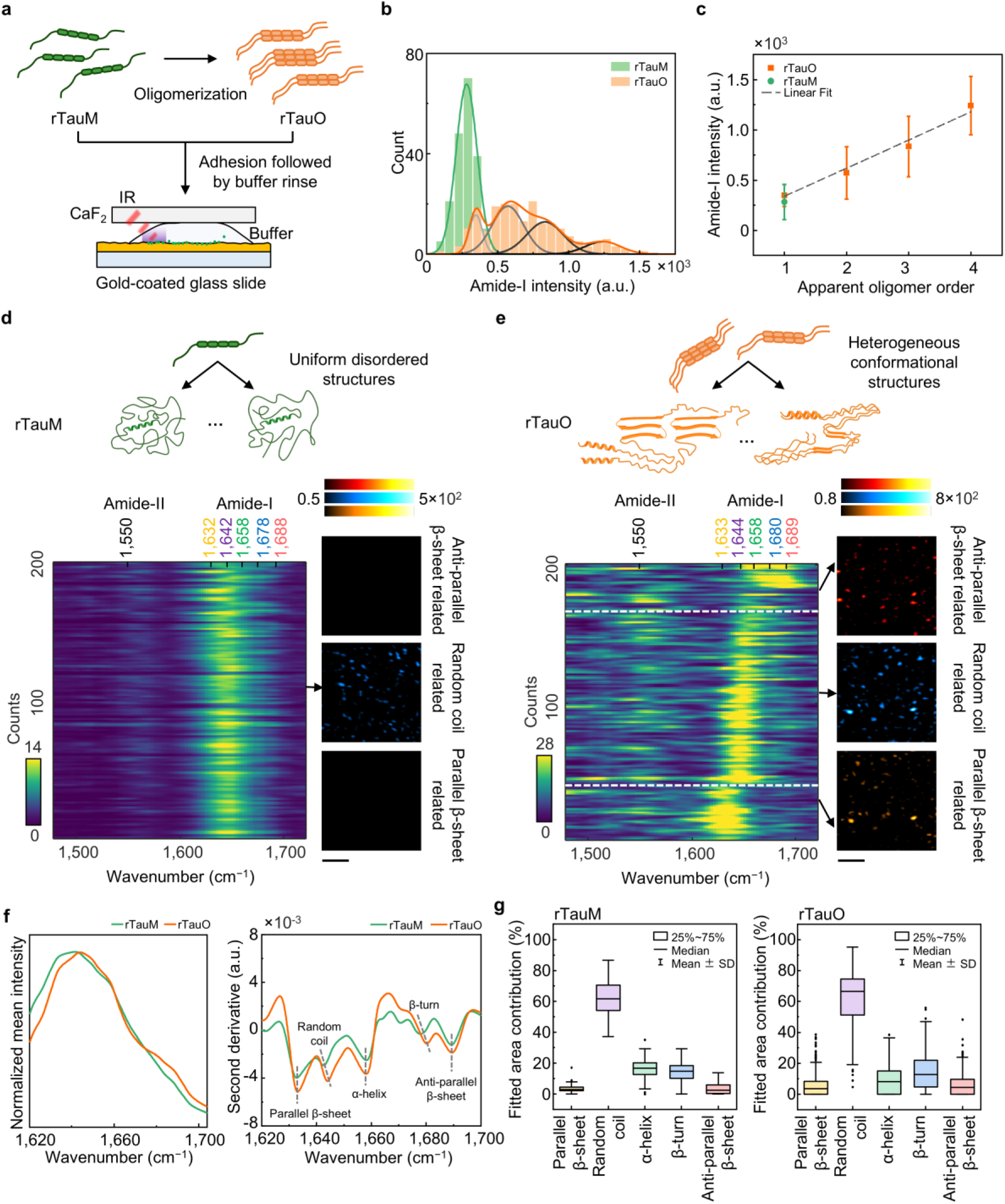
Conformational mapping of recombinant human 2N4R tau proteins. a, Schematic of oligomerization from recombinant human 2N4R human tau monomers (rTauM) to oligomers (rTauO) and imaging workflow. **b,** Amide-I band intensity histograms of rTauM (green) and rTauO (orange) particles with Gaussian fits (*n* = 200 for each). **c,** Amide-I band intensity versus apparent oligomer order (rTauM, green; rTauO, orange). Order 1 corresponds to the rTauM population. Data points represent the mean of each peak in **b**. Error bars indicate the fitted Gaussian FWHM. **d–e,** Heatmaps of IR-AMES spectra from monomers (**d**) and oligomers (**e**). Top, schematic illustrations of predominantly disordered rTauM and structurally heterogeneous rTauO assemblies. Detailed conformations predicted by AlphaFold3 are provided in the **Supplementary Note 6** and **Supplementary** Fig. 18. Bottom left, heatmaps of single-particle IR-AMES spectra. Bottom right, representative structure-selective IR-AMES images at bands associated with antiparallel β-sheet, random coil and parallel β-sheet structures. Scale bars: 2 µm. **f,** Mean normalized amide-I spectra of rTauM and rTauO (left) and the corresponding second-derivative spectra (right). Dashed lines indicate the peak minima used to define the component centers. **g,** Single-particle distributions of fitted secondary-structure contributions for rTauM and rTauO.

In contrast, rTauO exhibits distinct spectral heterogeneity (**Fig. 3e**). Most oligomers remain dominated by random-coil features near 1,644 cm^−1^, whereas smaller subsets show enhanced parallel β- sheet or antiparallel β-sheet signatures near 1,633 and 1,689 cm^−1^, respectively, consistent with the structure-selective images. This spectral diversity indicates that early tau oligomerization generates structurally heterogeneous assemblies rather than a single, well-defined conformation. Consistently, AlphaFold3 predictions of tau dimers and trimers frequently reveal β-sheet-rich motifs embedded within otherwise disordered conformations (**Supplementary Fig. 18b**). Bulk FTIR confirmed the protein-rich composition of both samples but obscured the underlying structural heterogeneity through ensemble averaging (**Supplementary Fig. 19**).

To quantify spectral heterogeneity at the single-particle level, we applied a conventional multi-component fitting approach^18, 32, 42^ to the amide-I spectra (**Supplementary Fig. 20**). Mean IR-AMES spectra and their second derivatives were used to identify characteristic spectral minima and guide component assignment (**Fig. 3f**). Fitting of individual spectra shows that rTauM is dominated by a relatively narrow distribution of random-coil-rich conformations, whereas rTauO exhibits broader structural distributions with increased contributions from parallel and antiparallel β-sheet components (**Fig. 3g**). Together, these results demonstrate that recombinant tau oligomerization is accompanied by the emergence of structurally heterogeneous, β-sheet-containing conformations, and establishes a foundation for IR-AMES to probe disease-relevant tau assemblies with single-particle chemical specificity.

## AD patient-derived tau oligomers are featured by antiparallel β-sheet and RNA enrichment

Disease-associated tau assemblies may display distinct structural and compositional signatures linked to neurotoxicity. To resolve these features at the single-particle level, we examined tau oligomers (TauO) and fibrils (TauF) isolated from postmortem AD brains^43, 44^. Western blot analysis confirmed their distinct assembly states (**Supplementary Fig. 21**). We next characterized their morphology. AFM images revealed that AD-derived tau oligomers (AD TauO) predominantly adopt spherical or ellipsoidal morphologies with heights of 5–8 nm and lateral dimensions of ∼40 nm, whereas AD-derived tau fibrils (AD TauF) form short, fragmented rod-like structures with increased length and height (**Fig. 4a** and **Supplementary Fig. 22**). We next assessed the neurotoxicity of these tau assemblies using iPSC-derived neurons. Neurons exposed to AD TauO exhibited significantly increased cytotoxicity, as indicated by elevated lactate dehydrogenase (LDH) release and enhanced cleaved caspase-3 signaling, compared with AD TauF (**Fig. 4b, c** and **Supplementary Fig. 23**). Age-matched control brain–derived tau oligomers (Ctrl TauO) and control tau fibrils (Ctrl TauF) showed lower toxicity than AD TauO. These results suggest that soluble tau oligomers derived from AD brains represent the most neurotoxic tau species.

**Fig. 4.**
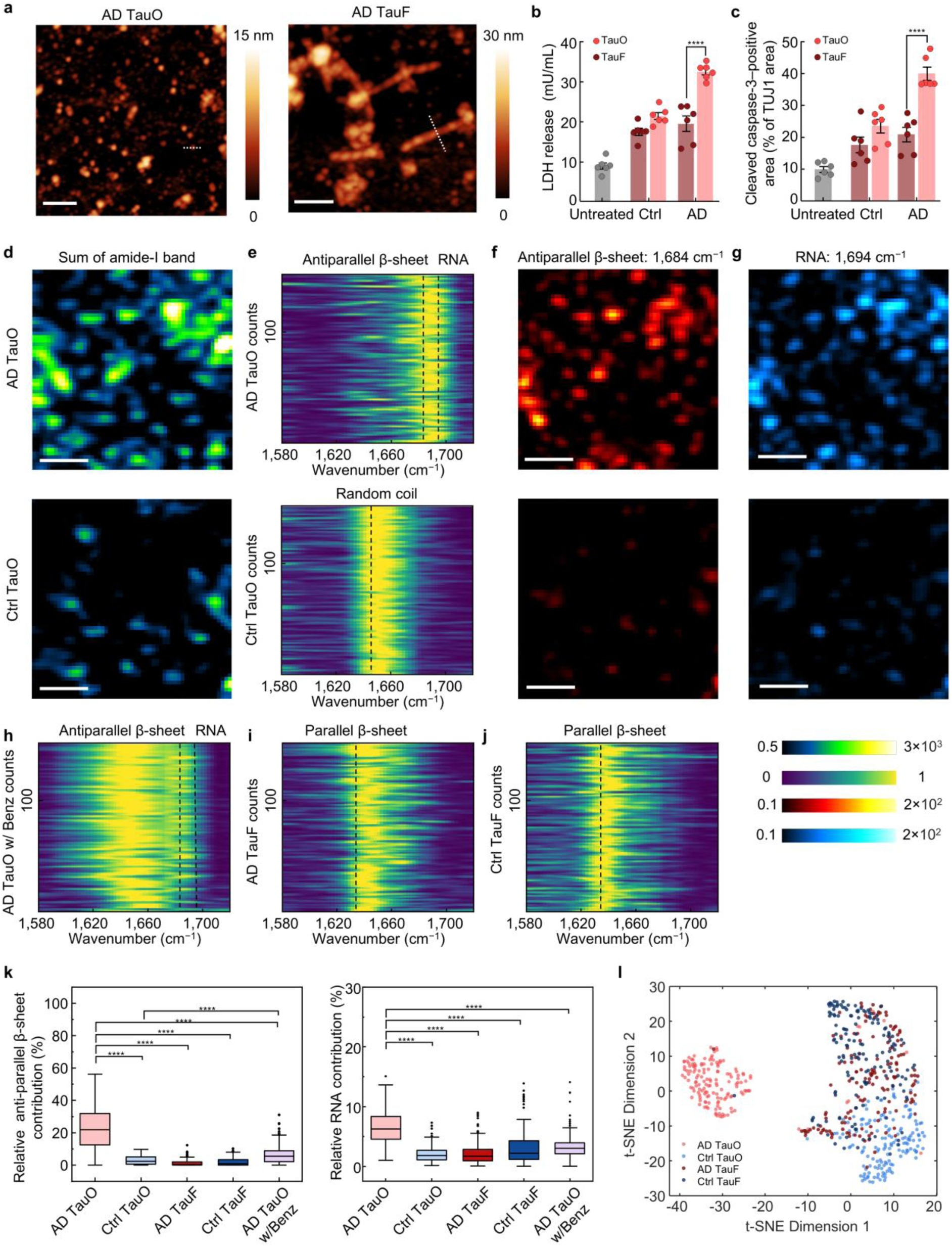
Structural and compositional mapping of human-derived tau oligomers provide insights into neuronal toxicity. a, Atomic force microscopy images of Alzheimer’s disease patient derived tau oligomers (AD TauO) and fibrils (AD TauF). Scale bars: 250 nm. AD TauO appear as spherical or ellipsoidal particles wih heights of 5–8 nm and lateral dimensions of ∼40 nm. AD TauF appear as short fragmented rods with heights of ∼10 nm, lateral widths of ∼30–50 nm, and lengths ranging from 100 to 500 nm. Dimensions were measured along the white dashed lines, details are provided in **Supplementary** Fig. 22. **b–c,** Cytotoxicity of iPSC-derived neurons treated with human tau for 24 h, quantified by LDH release (**b**) and cleaved caspase-3–positive area relative to TUJ1 (**c**). *n* = 6. Data were expressed as mean ± s.d. Column means were compared using two-way ANOVA, with ****p < 0.0001. **d,** IR-AMES image of AD TauO and age-matched normal human derived tau oligomers (Ctrl TauO) at the amide-I band. Scale bars: 1 µm. **e,** Heatmaps of IR-AMES spectra from AD TauO and Ctrl TauO, *n* = 150. Spectra were normalized to 0–1. **f–g,** IR-AMES image of AD TauO and Ctrl TauO at the antiparallel β-sheet channel (**f**) and RNA channel (**g**). Scale bars: 1 µm. **h–j,** Heatmaps of IR-MAES spectra from AD TauO with endonuclease benzonase (AD TauO w/Benz) treatment (**h**), AD TauF (**i**) and normal human derived tau fibrils (Ctrl TauF) (**j**), *n* = 150. Spectra were normalized to 0–1. **k,** Quantification of antiparallel β- sheets and RNA content from human tau in **e** and **h–j**. All groups were expressed as mean ± s.d. Column means were compared using one-way ANOVA, with ****p < 0.0001, and ns for not significance. **l,** t-SNE visualization of all spectra from individual tau assemblies, revealing structure-dependent clustering patterns. Each dot indicates a single-particle spectrum.

To identify molecular features associated with AD tau oligomer toxicity, we applied IR-AMES to characterize the structure and composition of individual tau assemblies. IR-AMES imaging at the amide- I band window revealed strong signals from both AD and Ctrl TauO (**Fig. 4d**). IR-AMES spectral heatmaps showed distinct heterogeneity in AD TauO, with major contributions near 1,684 cm^−1^ and 1,694 cm^−1^, corresponding to antiparallel β-sheet and RNA-associated (uracil C=O) vibrational modes, respectively. In contrast, spectra of Ctrl TauO derived from non-AD brains were dominated by a peak near 1,646 cm^−1^, indicating random-coil conformations (**Fig. 4e**). Consistent with these observations, IR- AMES images acquired at the antiparallel β-sheet (**Fig. 4f**) and RNA (**Fig. 4g**) channels further revealed strong enrichment of both signals in AD TauO compared with Ctrl TauO. To validate the RNA contribution, AD TauO were treated with endonuclease benzonase prior to imaging. This enzymatic RNA digestion remarkably reduced the ∼1,694 cm^−1^ RNA-associated peak, accompanied by broadening of the amide-I features (**Fig. 4h**), indicating that RNA binding contributes to the antiparallel β-sheet. In comparison, both AD- and control-derived TauF exhibited dominant features near ∼1,630 cm^−1^, corresponding to parallel β-sheet structures, with minimal RNA-associated signal (**Fig. 4i**, **j** and **Extended Data Fig. 4**). Further AFM and IR-AMES analysis showed that AD TauF predominantly consisted of short fibrillar structures, which may reflect fragmentation during sample isolation and preparation (**Supplementary Fig. 24**).

Quantitative analysis based on multi-component fitting confirmed these signatures (**Fig. 4k** and **Extended Data Fig. 5**). AD TauO exhibits significant enrichment of both antiparallel β-sheet and RNA compared with all control groups. Benzonase treatment reduced RNA signals and was accompanied by a partial decrease in antiparallel β-sheet signal. This decrease may reflect partial spectral overlap between RNA-associated and antiparallel β-sheet bands or a possible influence of RNA association on oligomer conformation, yet the residual antiparallel β-sheet content remained significantly higher than in control tau species. Collectively, IR-AMES measurements reveal that tau oligomers from human AD brains are distinguished by the coexistence of antiparallel β-sheet structures and RNA enrichment. Full fingerprint- region FTIR spectra further confirm that protein is the dominant component in the brain-derived tau preparations (**Supplementary Fig. 25**). However, ensemble measurements predominantly resolve the parallel β-sheet signatures at 1,630 cm^−1^ in fibrillar assemblies, whereas the oligomer-specific antiparallel β-sheet and RNA presence are obscured, showing the significance of single particle analysis.

To ensure that these structural distinctions arise independently of peak fitting, we applied an unsupervised dimensionality reduction algorithm, t-distributed stochastic neighbor embedding (t-SNE), to the single-particle amide-I spectra. As shown in **Fig. 4l**, t-SNE allowed spectral clustering without predefined structural assignments. AD TauF and Ctrl TauF largely overlapped in the embedding space, consistent with their parallel β-sheet structures^5^. In contrast, AD TauO formed a clearly separated cluster, whereas Ctrl TauO occupied a distinct but partially proximal region relative to fibrillar assemblies. Centroid-distance analysis further confirmed the close clustering of fibrillar species and the distinct separation of AD TauO from other tau assemblies (**Supplementary Fig. 26**). Together, these findings establish a connection between structure, composition and neurotoxicity of AD TauO.

To assess the applicability of IR-AMES beyond tau, we further measured in vitro–formed α- synuclein amyloid fibrils prepared and characterized as described previously^45^. IR-AMES resolved micrometre-scale fibrillar structures and amide-I spectra dominated by the parallel β-sheet band near 1,630 cm^−1^, consistent with the parallel cross-β architecture of this fold^45^ and with ensemble FTIR measurements (**Extended Data Fig. 6**). These results provide an independent, structurally defined amyloid benchmark and support the application of IR-AMES to protein assemblies beyond tau.

## IR-AMES reveals a strong interaction between AD TauO and lipid nanodiscs

Given the strong implication of tau–membrane interactions in neurotoxicity and membrane disruption^46–49^, we examined how these molecular signatures identified in AD-derived TauO influence tau–lipid interaction using lipid nanodiscs (NDs) as a testbed with defined compositions. The nanodiscs were assembled using an amphipathic α-helical 18A scaffold peptide modified with hexanoic acid^50^, providing a chemically defined membrane environment (**Fig. 5a**). Nanodiscs composed of phosphatidylcholine (PC) alone or PC mixed with negatively charged phosphatidylserine (PS) were prepared in buffer and validated by size measurements and FTIR (**Supplementary Fig. 27**). Nanodiscs alone under IR-AMES showed co-localized amide-I and lipid signals, with spectra dominated by α-helix and lipid features, consistent with a homogeneous scaffold protein and lipid bilayer architecture (**Figs. 5b–d**).

**Fig. 5.**
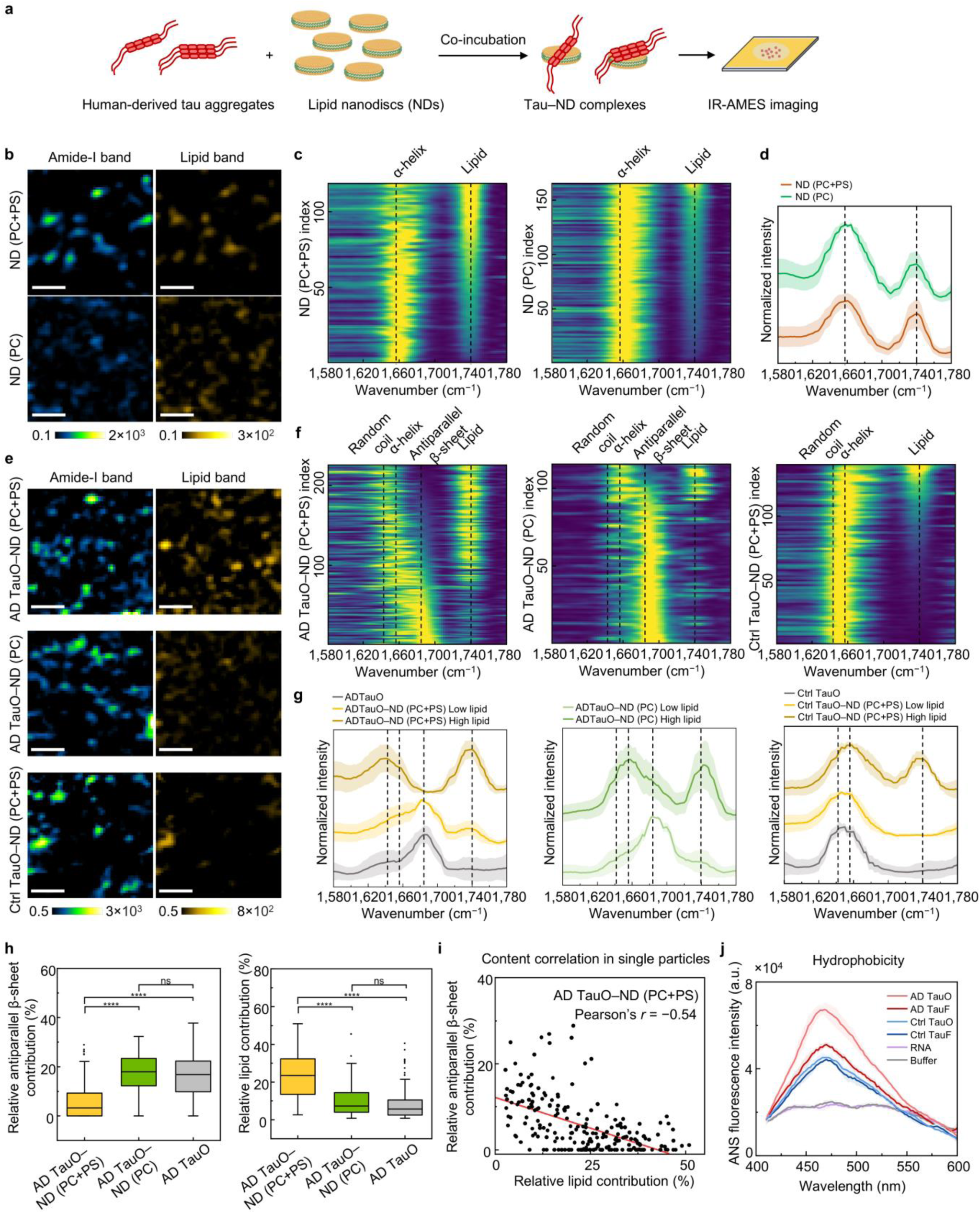
IR-AMES investigation of human-derived tau oligomer and lipid membrane interactions. a, Schematic illustrating the co-incubation of human-derived tau aggregates with lipid nanodiscs (NDs) to form tau–ND complexes for IR-AMES imaging. **b,** Representative IR-AMES images of NDs composed of phosphatidylcholine and phosphatidylserine (PC+PS) or PC only, shown for integrated amide-I and lipid signals. Scale bars, 1 µm. **c,** Heatmaps of IR-AMES spectra from ND (PC+PS) (*n* = 119) and ND (PC) (*n* = 165). Spectra were normalized to 0–1 and ordered by lipid intensity. **d,** Average spectra of NDs. Solid lines: mean spectra, shaded regions: standard deviation. **e,** Representative images of AD TauO and Ctrl TauO following NDs incubation. Scale bars, 1 µm. **f,** Heatmaps of spectra from AD TauO–ND (PC+PS) (*n* = 225), AD TauO–ND (PC) (*n* = 115), and Ctrl TauO–ND (PC+PS) (*n* = 140), highlighting distinct protein secondary-structure and lipid-associated spectral features. Spectra for AD TauO–ND spectra were ordered by antiparallel β-sheet contribution. **g,** Average spectra of lipid-poor (*n* = 25) and lipid-enriched (*n* = 25) subsets derived from **f**, compared with tau aggregates alone (*n* = 150 for each). Spectra were normalized to 0–1 and vertically offset for display in **d** and **g**. **h,** Quantification of antiparallel β-sheet and lipid contributions from IR-AMES spectra. All groups were expressed as mean ± s.d. Column means were compared using one-way ANOVA, with ****p < 0.0001, and ns for not significance. **i,** Single-particle correlation analysis of lipid content and antiparallel β-sheet contribution in AD TauO–ND (PC+PS), revealing a moderate negative correlation (Pearson’s *r* = –0.54). **j,** 8-anilino- 1-naphthalenesulfonic acid (ANS) fluorescence spectra of human tau (*n* = 3, solid lines: mean spectra, shaded regions: standard deviation), indicating enhanced hydrophobicity of AD TauO relative to controls.

Upon co-incubation of tau assemblies with nanodiscs, AD TauO exhibited distinct interaction with the PS-containing membranes (**Figs. 5e-g**). Spectra of AD TauO–ND (PC+PS) complexes resolved into two subpopulations (**Fig. 5f, g**; left column): lipid-poor particles retained dominant antiparallel β-sheet signatures characteristic of AD TauO alone, whereas lipid-enriched particles exhibited reduced antiparallel β-sheet contributions together with increased disordered and α-helical spectral features. The latter likely reflects a combination of structural remodeling and contributions from the α-helical nanodisc scaffold. In comparison, this membrane-induced structural shift was significantly attenuated in neutral ND (PC) conditions (**Fig. 5f, g**; middle column), and largely absent for Ctrl TauO (**Fig. 5f, g**; right column), which retained predominantly random-coil signatures. Together, these data suggest a selective interaction between AD TauO and negatively charged lipids.

Quantitative analysis confirmed that AD TauO–ND (PC+PS) complexes exhibited increased lipid content accompanied by reduced antiparallel β-sheet component relative to the AD TauO alone or ND (PC) conditions (**Supplementary Fig. 28** and **Fig. 5h**). Importantly, single-particle correlation analysis revealed a moderate negative correlation between lipid and antiparallel β-sheet contribution (Pearson’s r = –0.56) in AD TauO–ND (PC+PS) complexes (**Fig. 5i**). This indicates that membrane interaction is accompanied by destabilization of β-sheet conformations. In contrast, AD TauF maintained dominant parallel β-sheet signatures with minimally detectable lipid signal (**Supplementary Fig. 29**), indicating limited fibril–membrane interaction and relatively weak photothermal contrast of small lipid membranes compared to the tau fibrils.

Consistent with the preferential membrane association of AD TauO, through a fluorescence assay^51^, we find stronger surface hydrophobicity of AD TauO relative to Ctrl TauO and fibrils (**Fig. 5j**). Together, these results indicate that AD TauO preferentially engages anionic membranes and such tau–membrane interaction is associated with partial loss of β-sheet structure. Collectively, these findings suggest a mechanistic connection between structure and pathogenic activity of tau oligomers via interactions with lipid membranes.

## Conclusions

Single-molecule imaging has revealed extensive molecular heterogeneity within biological systems. Yet, current approaches largely rely on either fluorescence labeling^52–55^ or scattering contrast^19–22^, both lacking direct structural or compositional information. IR-AMES fills this gap by photothermally encoding IR spectral information into scattering contrast. By combining a surface-confined evanescent field with an orthogonal detection geometry, IR-AMES enables label-free, widefield IR spectroscopic imaging with single-protein sensitivity and nanoscale resolution. With multi-component spectral analysis, it allows direct structural and compositional profiling of individual biomolecular assemblies in buffer, where existing structural and spectroscopic approaches remain limited.

Applying IR-AMES to tau oligomers reveals a link between nanoscale chemical feature and pathogenic function. During recombinant tau aggregation, the emergence of β-sheet structures and increasing conformational diversity from initially disordered monomers are observed. In human-derived samples, AD tau oligomers are distinguished by enrichment of antiparallel β-sheets and RNA, accompanied by increased hydrophobicity and neuronal toxicity. Cryo-electron microscopy of tau filaments from human tauopathies has revealed non-proteinaceous densities embedded within positively charged cavities, consistent with polyanionic cofactors that stabilize disease-specific folds^56^. Although the molecular identity of these cofactors remains unresolved, the RNA enrichment observed here suggests that nucleic acids may participate at early stages of tau assembly, prior to the emergence of mature fibrillar architectures. This interpretation is consistent with cellular evidence that tau oligomerization promotes association with RNA-binding proteins and N6-methyladenosine–modified RNA transcripts^43^.

Using lipid nanodiscs as a defined membrane mimic, we found that AD tau oligomers preferentially associate with anionic lipid membranes. This membrane engagement is accompanied by destabilization of antiparallel β-sheet–rich conformations and a shift toward less ordered structural states, indicating that tau oligomers are conformationally labile and sensitive to local microenvironments. The preferential interaction between AD tau oligomers and negatively charged lipid nanodiscs further suggests selective engagement of AD tau oligomers with negatively charged membrane interfaces, including the outer nuclear membrane. In consistence, tau oligomers have also been reported to localize to and perturb the nuclear envelope, inducing membrane invagination and lamina disruption^49^.

In summary, IR-AMES establishes a nanoscale spectroscopic imaging platform for ultrasensitive fingerprinting of biomolecular assemblies under native conditions. This capability opens opportunities in viral subtyping, biopharmaceutical quality control, extracellular vesicle profiling, and microbial identification. More broadly, IR-AMES heralds exciting potential for uncovering structure–function relationships in complex biomolecular systems^57^, from understanding lipid–protein interactions to screening small compounds that remodel pathological protein conformations and modulate disease- associated activity.

## Methods

### Materials

Gold-coated glass slides with gold thickness of ∼50 nm were purchased from Biosensing Instrument. CaF_2_ substrates (CAFP13-0.2R) were purchased from Crystran. PMMA nanoparticles (MMA500, MMA100, MMA75, MMA50) were purchased from Phosphorex. Amino PMMA nanoparticles (500 nm) were purchased from Polysciences (07763-5). Fluorescent PS nanoparticles were purchased from Bangs Laboratories (FSDG001). IgM from human serum (I8260), myoglobin (M0630), concanavalin A (Con A, C2010), DL-dithiothreitol (DTT, D9779), arachidonic acid (AA, A3611), benzonase (Benz, E1014), 8-Anilino-1-naphthalenesulfonic acid ammonium salt (ANS, 10417), fluorescein isothiocyanate isomer I (FITC, F7250), sodium chloride (S9888), and zinc chloride (208086) were purchased from Sigma–Aldrich. Recombinant human tau protein was purchased from R&D Systems, Inc (SP-495). Universal human reference RNA (QS0639) and Tris (1 M, pH 8.0, AM9855G) were purchased from Thermo Fisher Scientific. Recombinant wild-type α-synuclein was purified and assembled into amyloid fibrils in Tris buffer as described previously^45^.

### Generation of human tau oligomers and fibrils

Anonymous post-mortem human prefrontal cortex (Brodmann area 10) tissues from age-matched control (*n* = 8) and Alzheimer’s disease (*n* = 8) cases were obtained from the Emory Goizueta Alzheimer’s Disease Research Center brain bank (NIH P30AG066511). All samples were fresh-frozen and de-identified. Tau oligomer (TauO) and fibril (TauF) fractions were prepared as previously described^43, 44^. Briefly, 100 mg frozen tissue was homogenized in 10 volumes of Hsiao TBS buffer (50 mM Tris, pH 8.0, 274 mM NaCl, 5 mM KCl) supplemented with protease and phosphatase inhibitors. Homogenates were centrifuged at 28,000 rpm (Beckman Coulter Optima MAX-TL ultracentrifuge, TLA-55 rotor) for 20 min at 4 °C. The supernatant (S1; TBS-soluble fraction) was further centrifuged at 55,000 rpm for 40 min at 4 °C. The resulting pellet (TauO fraction) was resuspended in TE buffer (10 mM Tris, 1 mM EDTA, pH 8.0; 4× volume relative to starting tissue weight), aliquoted, and stored at −80 °C.

The initial pellet (P1) was homogenized in buffer B (10 mM Tris, pH 7.4, 800 mM NaCl, 10% sucrose, 1 mM EGTA, 1 mM PMSF; ∼5× tissue wet weight) and centrifuged at 22,000 rpm for 20 min at 4 °C. The supernatant (S2) was incubated with 1% Sarkosyl at 37 °C for 1 h with rotation and centrifuged at 55,000 rpm for 1 h at 4 °C. The sarkosyl-insoluble pellet (P3; TauF fraction) was resuspended in 50 µL TE buffer and stored at −80 °C.

Tau species in TauO and TauF fractions were analyzed by native PAGE. Total tau levels were quantified by immunoblot using 3–12% SDS–PAGE and the Tau-5 antibody, with recombinant tau standards for calibration. Fractions were normalized to 20 µg mL⁻¹ total tau for storage and downstream applications.

The human brain tissue samples used in this study were all de-identified. All studies included both sexes, and results were integrated by covariate analysis. Donor characteristics are summarized in **Supplementary Table 3**.

### Preparation of lipid nanodiscs

Nanodiscs were assembled by incubating liposomes composed of 1,2- dioleoyl-sn-glycero-3-phosphocholine (DOPC) or 70 mol% DOPC and 30 mol% 1,2-dioleoyl-sn-glycero-3-phospho-L-serine (DOPS) with the scaffold peptide Hex18A^50^ at a ratio of 1/30 at 4°C with gentle shaking for 16 h. The mixture was purified by size-exclusion chromatography using a Superose 6 10/300 column equilibrated in 25 mM Tris-HCl, 100 mM NaCl. Fractions containing nanodiscs were pooled together and concentrated to 100 µM before use.

### IR-AMES setup

The IR-AMES system (**Supplementary Fig. 4**) was built on an inverted microscope frame (IX70, Olympus) and integrated a TIR visible probe with widefield mid-IR photothermal excitation. The visible probe light was provided by a lab-built nanosecond pulsed 450-nm laser (K450F03FN-2.6W, Precision Micro-Optics Inc.) (**Supplementary Fig. 4**, bottom). The beam was collimated using an achromatic doublet (AC254-030-A-ML, Thorlabs) and then focused onto the back focal plane of an oil-immersion objective (UPLAPO60XOHR, 60×, NA 1.50, Olympus) by a second lens (AC508-180-A-ML). The TIR incidence angle was tuned by translating the fiber output of the laser. An evanescent field was generated above the interface to excite scattering from nano-objects on the surface. The scattered light was collected by the same objective and separated from the illumination path using a beam splitter (BSW10R, Thorlabs). A spatial barrier placed near the back focal plane blocked the specular reflection, allowing only scattered light to reach a CMOS camera (BFS-U3-20S4M-C, FLIR) for image acquisition. The mid-IR pump light was provided by a pulsed QCL laser (MIRcat 2400, Daylight Solutions), tunable from 900 to 1,800 cm^−1^. The IR beam was expanded fivefold and relayed using a pair of concave mirrors (CM254-50-P01 and CM254-250-P01, Thorlabs) (**Supplementary Fig. 4**, top). The expended IR beam was then weakly focused onto the sample at 45° incidence in *p* polarization using an off-axis parabolic mirror (MPD124-P01, Thorlabs). The mid-IR power at the sample plane was measured using a power meter (PM16-401, Thorlabs) for spectra normalization.

The system was electrically controlled by a four-channel delay pulse generator (9254, Quantum Composers), which provided a 100-kHz master clock to synchronize the pump pulses, probe pulses, and camera exposure (**Supplementary Fig. 5**). The mid-IR pulse train was set at a 50% duty cycle, with pulse widths of 1 µs for measurements in air and 40–80 ns for measurements in aqueous environments (80 ns for PMMA beads, 40 ns for biomolecules). The camera was triggered to recorded IR-on and IR- off frames with the same exposure time (e.g., 5 ms, corresponding to 500 IR pulses per frame) sequentially, and IR-encoded scattering images were generated by two-frame subtraction at each wavenumber. For hyperspectral imaging, the QCL operated in a “Step-and-Measure” mode, sending a start-of-scan trigger at each wavenumber that initiated a synchronized pump–probe–camera acquisition sequence. This acquisition cycle was repeated at every spectral step until the scan reached the final wavenumber, with a dead interval of ∼250 ms between adjacent steps. The estimated power density at the sample plane was ∼0.1–6 kW/cm^2^ for the visible probe and ∼0.05–1.5 kW/cm^2^ for the mid-IR pump, depending on pulse width and effective illumination area. The probe power density and camera exposure time were optimized for each measurement (**Supplementary Table 4**).

### Sample preparation

For PMMA nanoparticle detection, PMMA beads were diluted 100-1000 times in deionized (DI) water and then spin-coated onto gold-coated glass slides, and dried in air. For measurements in aqueous environments, the dried PMMA-coated substrates were gently rinsed with DI water to remove loosely attached particles. A small volume of DI water (<0.1 µL) was added onto the surface and sealed with a CaF_2_ coverslip to form a thin aqueous layer (<150 nm at the edge) for IR- AMES imaging. For IgM detection in air, IgM was diluted to 1 nM in PBS and incubated on a cleaned gold-coated coverslip for >1 h at 4℃ to allow unspecific binding. The surface was gently rinsed with PBS and DI water to remove unbound molecules and residual salts, then dried in air before IR-AMES imaging. For IgM detection in solution, IgM was prepared and adsorbed as above, followed by a gentle PBS rinse. A small volume of PBS (<0.1 µL) was added and sealed with a CaF_2_ coverslip to form a thin aqueous layer (<150 nm at the edge) for IR-AMES measurements. Tau assemblies, nanodiscs, and tau– nanodisc complexes were immobilized on gold substrates via nonspecific adsorption to minimize perturbation of native structures. Incubation times were adjusted according to sample concentration and adsorption kinetics until sufficient individual particles were stably adsorbed on the surface. Control experiments at two different incubation times excluding motion-induced artifacts in the solution condition are shown in **Supplementary Fig. 30**. Detailed procedures are described in **Supplementary Note 7**.

### Data processing

IR-AMES imaging data were acquired with custom MATLAB scripts and analyzed with ImageJ. Data plotting and statistical analysis were performed in Origin and MATLAB. Pseudocolor was added to the IR-AMES and fluorescence images with ImageJ. Spectral analysis and multi-component fitting of single-particle spectra were performed in MATLAB. Detailed image and spectral analysis workflows are described in **Supplementary Note 8**. Biomolecule illustrations were created with BioRender. All values labeled “a.u.” in the figures represent arbitrary units.

### FTIR measurement

The FTIR spectra were acquired using an attenuated total reflection FTIR spectrometer (Nicolet Nexus 670, Thermo Fisher Scientific). The spectra resolution is 2 cm^−1^ and each spectrum was measured with 128 scanning. Baseline correction and spectral normalization were applied using the instrument software.

### AFM imaging

All AFM experiments were carried out in air using AC/tapping mode with a soft tapping mode tip (2 N/m) (240AC-NA-10, Nanoandmore USA) with the scan rate of 1.0 Hz with with the Asylum AFM Cypher S/ES instrument.

### Neuronal toxicity assay

Human induced pluripotent stem cells (iPSCs) were obtained from the JAX iPSC Collection (JIPSC001000). Neural progenitor cells (NPCs) were generated using STEMdiff™ Forebrain Neuron Differentiation Medium (Cat# 08600) on Matrigel-coated plates and maintained in STEMdiff™ Neural Progenitor Medium 2 (Cat#08560) according to the manufacturer’s instructions. For terminal differentiation, passage-matched NPCs were transduced with NEUROG2 lentivirus (MOI 3; GeneCopoeia, LPP-T7381-Lv105-A00-S). Medium was replaced 24 h post-transduction to remove residual virus, generating induced neurons (iNeurons). At 6 days after NGN2 derived neuronal differentiation, iNeurons were treated with 40 ng ml⁻¹ TauO or TauF fractions extracted from post-mortem Alzheimer’s disease or age-matched control brain tissue for 24 h. Conditioned medium was collected for lactate dehydrogenase (LDH) release assays using the CyQUANT™ LDH Cytotoxicity Assay (Thermo Fisher Scientific, C20301) according to the manufacturer’s protocol. Absorbance at 490 nm and 680 nm was measured using an Epoch microplate spectrophotometer (BioTek). For morphological and apoptosis analyses, neurons were fixed in 4% paraformaldehyde 96 h after tau exposure and subjected to immunofluorescence staining for neuronal markers and cleaved caspase-3. Experimental procedures, assay conditions, and quantitative analysis are described in **Supplementary Note 9**.

### Hydrophobicity measurement

Hydrophobicity was quantified using ANS fluorescence on a SpectraMax i3x microplate reader (excitation: 380 nm; PMT gain: high; 50 flashes/read). Samples contained 15 µM ANS mixed with either 1.6 ng/µL human Tau or 40 ng/µL RNA in 20 mM HEPES buffer (150 mM NaCl). Mixtures were incubated for 15 min at room temperature and measured in 50-µL volumes in a 96-well plate.

### Statistics analysis

Statistical analysis was performed using Origin 2018. Data are presented as mean ± s.d. For experiments with factorial designs, group means were compared using two-way ANOVA followed by Bonferroni multiple-comparisons tests. For comparisons among independent groups without a complete factorial design, one-way ANOVA followed by Bonferroni multiple-comparisons tests was used. For single-particle imaging experiments, the sample size refers to the number of counted particles (>100 per group). For toxicity assays, the sample size was ≥6 per group. Statistical significance was defined as ***p < 0.001, ****p < 0.0001, and ns, not significant.

## Data availability

The data that support the findings of this study are included in the article and its Supplementary Information. Source data are provided with this paper.

## Acknowledgements

AFM imaging was performed in part at the Harvard University Center for Nanoscale Systems, a member of the National Nanotechnology Coordinated Infrastructure Network, which is supported by the National Science Foundation under NSF Award No. ECCS-2025158. The authors thank Jason Tresback for training and assistance with AFM imaging. The authors thank Haonan Zong, Zhongyue Guo and Jian Zhao for helpful discussions on the development of the IR-AMES system, Hongjian He and Guangrui Ding for help on AlphaFold3 predictions, and Pintian Lv for helpful discussions on IgM imaging.

## Author contributions

Q.X. and J.-X.C. conceived the concept. L.J. provided guidance on experiments related to tau preparation and biological interpretation. H.B. provided the nanodisc samples and contributed to biological interpretation. Q.X., J.-X.C., L.J. and H.B. designed the experiments. Q.X. performed the experiments and analyzed the data. Q.W. carried out neuronal toxicity assays D.J. developed the simulation models for scattering modulation. D.D. and M.L. contributed to the development of the lab-built visible laser and the IR-AMES system. E.S. prepared the human-derived tau samples. J.A. assisted with data analysis. Q.R. prepared the nanodisc samples. P.W.-S. and F.B. provided the α-synuclein amyloid fibrils, R.B. helped with the sample preparation. Q.X. and J.-X.C. wrote the manuscript with edits and approval from all authors.

---

**Extended Data Fig. 1.**
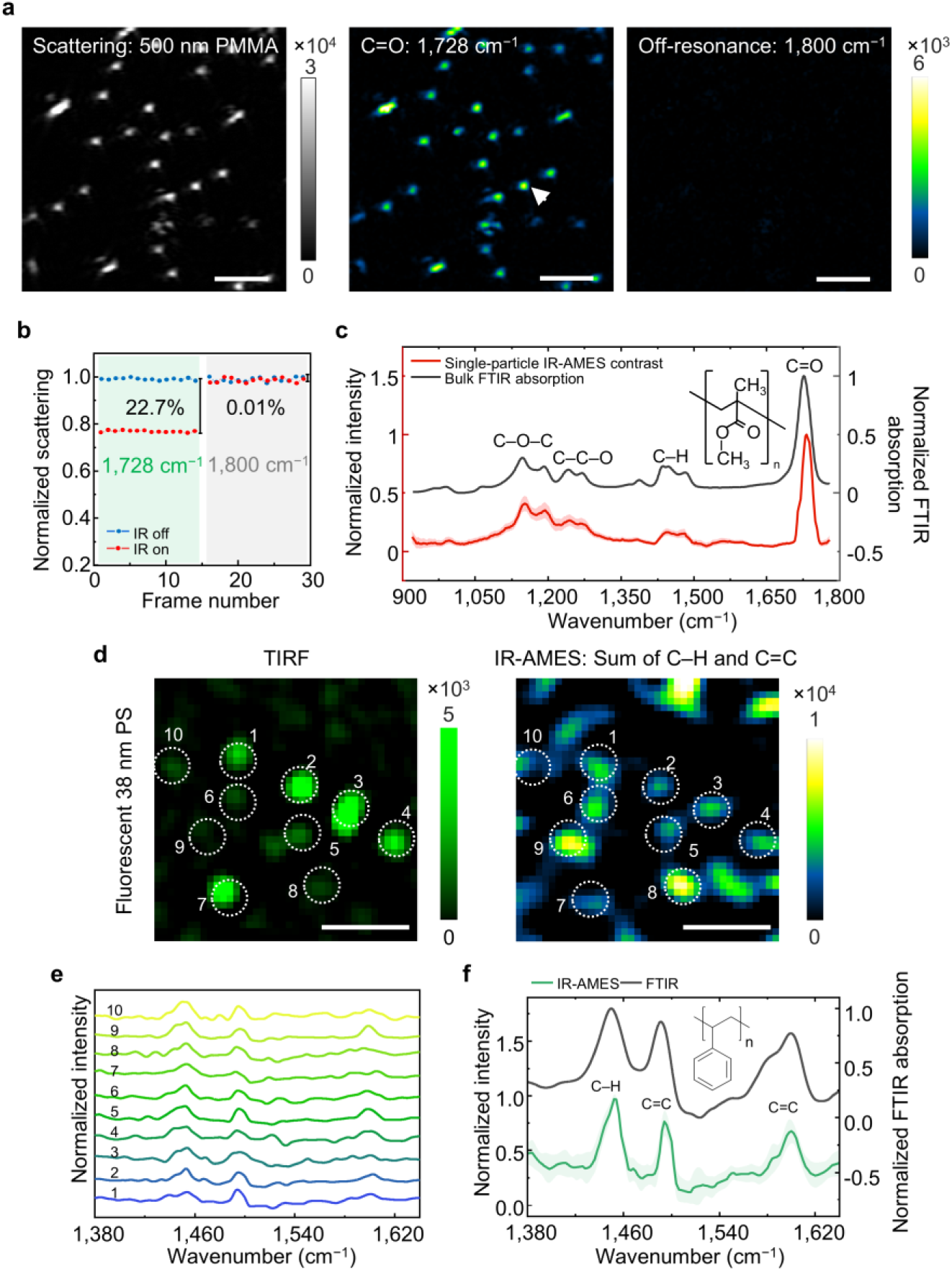
Experimental validation of enhanced scattering modulation and spectral fidelity in IR-AMES. a, Evanescent scattering image of single 500-nm PMMA particles at the IR-off state, and corresponding IR-AMES images at on-resonance (1,728 cm^−1^, C=O band) and off-resonance (1,800 cm^−1^). Scale bars: 3 µm. **b,** Scattering modulation at on- (1,728 cm^−1^) and off-resonance (1,800 cm^−1^) of the PMMA particle indicated in **a**. **c,** Spectral fidelity of IR-AMES measured from 500-nm PMMA beads (*n* = 50). Black line: FTIR spectrum, red line: mean IR-AMES spectra, shaded region: standard deviation. Inset: molecular structure of PMMA. **d,** Correlative TIRF and IR-AMES imaging of fluorescent 38-nm PS particles. Differences in relative particle intensities between the two modalities may reflect variations in fluorophore loading and their distinct contrast mechanisms. Scale bars: 1 µm. **e,** Normalized single-particle IR-AMES spectra of the ten numbered particles in **d**. **f,** Comparison of the mean IR-AMES spectrum of the ten PS particles with the bulk FTIR spectrum. Solid lines: mean spectra, shaded regions: standard deviation. Inset: molecular structure of PS.

**Extended Data Fig. 2.**
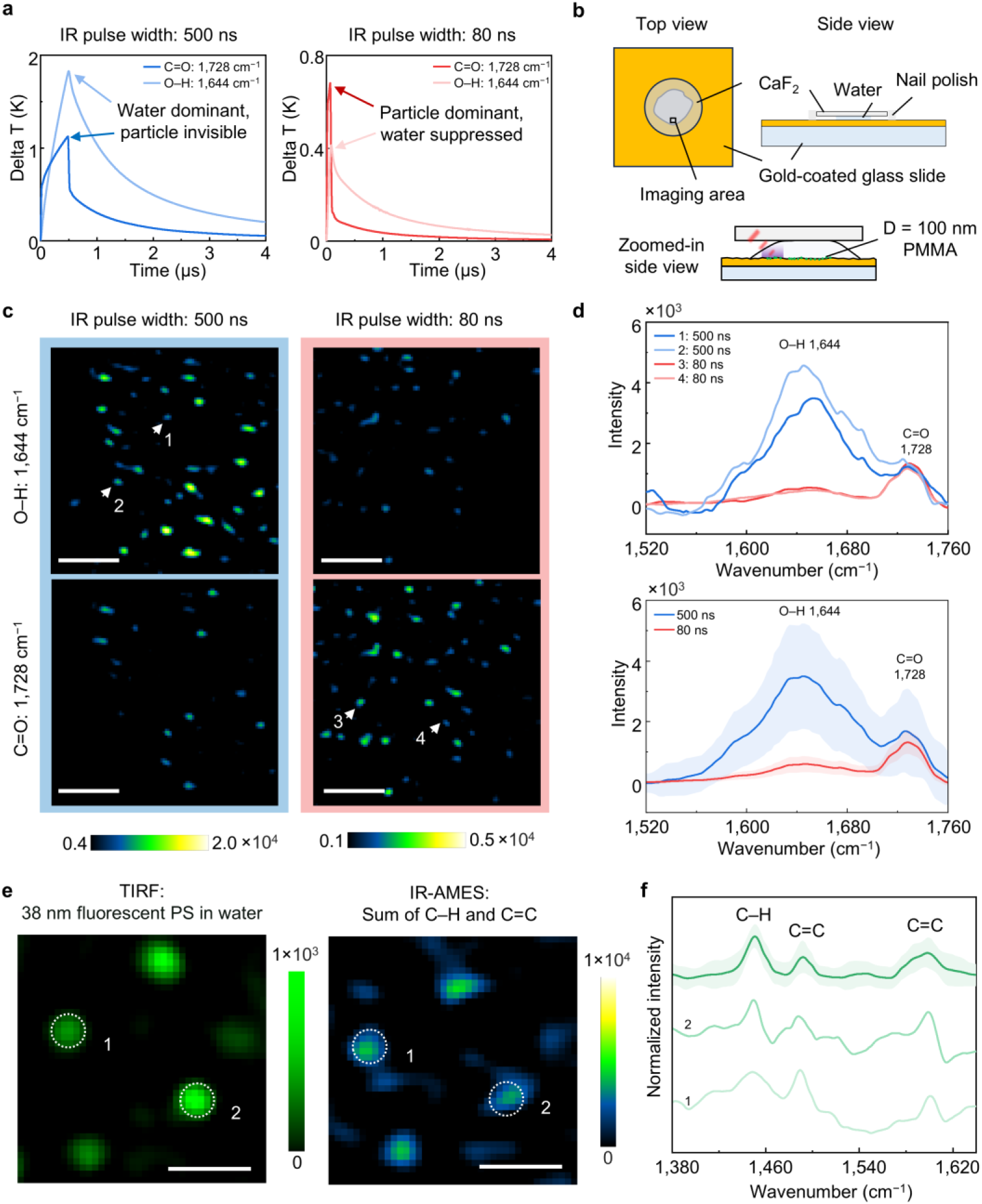
Water-suppression strategy enables IR-AMES detection of nanoparticles in water. a, Simulated thermal decay of a 100-nm PMMA nanoparticle in water. A long IR pulse (500 ns) produces strong water absorption at the O–H band (1,644 cm^−1^), making the PMMA C=O band (1,728 cm^−1^) invisible. A short 80-ns pulse confines heating to the particle, suppresses water absorption, and makes the bead signal dominant. **b,** Schematic of IR-AMES imaging in liquid environment with thin water layer and oblique IR incidence at the edge. **c,** IR-AMES images of 95-nm PMMA nanoparticles in water with 500-ns and 80-ns IR pulses. The average mid-IR power at the sample plane was 11.8 mW (500 ns) and 2.7 mW (80 ns) at 1,728 cm^−1^, and 15.5 mW (500 ns) and 3.5 mW (80 ns) at 1,644 cm^−1^. Scale bars: 1 µm. **d,** Representative single-particle IR-AMES spectra (top) and mean spectra (bottom) of 95-nm PMMA particles showing that under 80-ns IR excitation, the water background is suppressed and the PMMA peak appears. Solid lines: mean spectra, shaded regions: standard deviation (500 ns: *n* = 60; 80 ns: *n* = 64). **e,** Correlative TIRF and IR-AMES imaging of 38-nm fluorescent PS particles in water. Scale bars: 1 µm. **f,** Representative single-particle IR-AMES spectra and the mean spectrum of the numbered particles in **e**. Solid lines: mean spectra, shaded regions: standard deviation.

**Extended Data Fig. 3.**
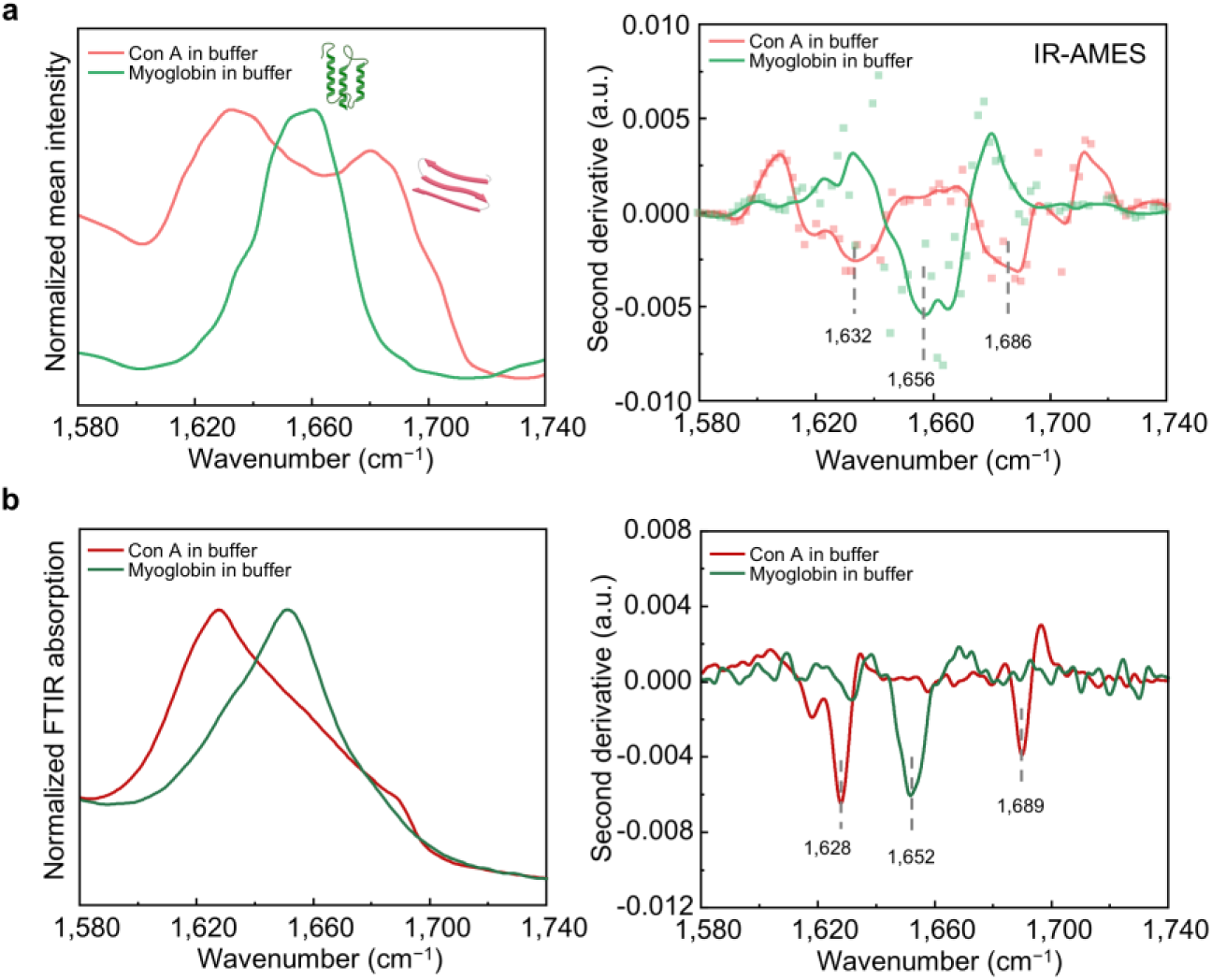
Validation of protein secondary-structure assignments using reference proteins. a, Mean IR-AMES spectra of concanavalin A (Con A) and myoglobin in buffer (left) and the corresponding second-derivative spectra (right). Dashed lines indicate characteristic minima associated with parallel β-sheet-rich Con A and α-helix-rich myoglobin. **b,** Corresponding bulk FTIR spectra (left) and second-derivative spectra (right). The agreement in spectral features and second-derivative minima supports the assignment of characteristic secondary-structure bands in the IR-AMES spectra.

**Extended Data Fig. 4.**
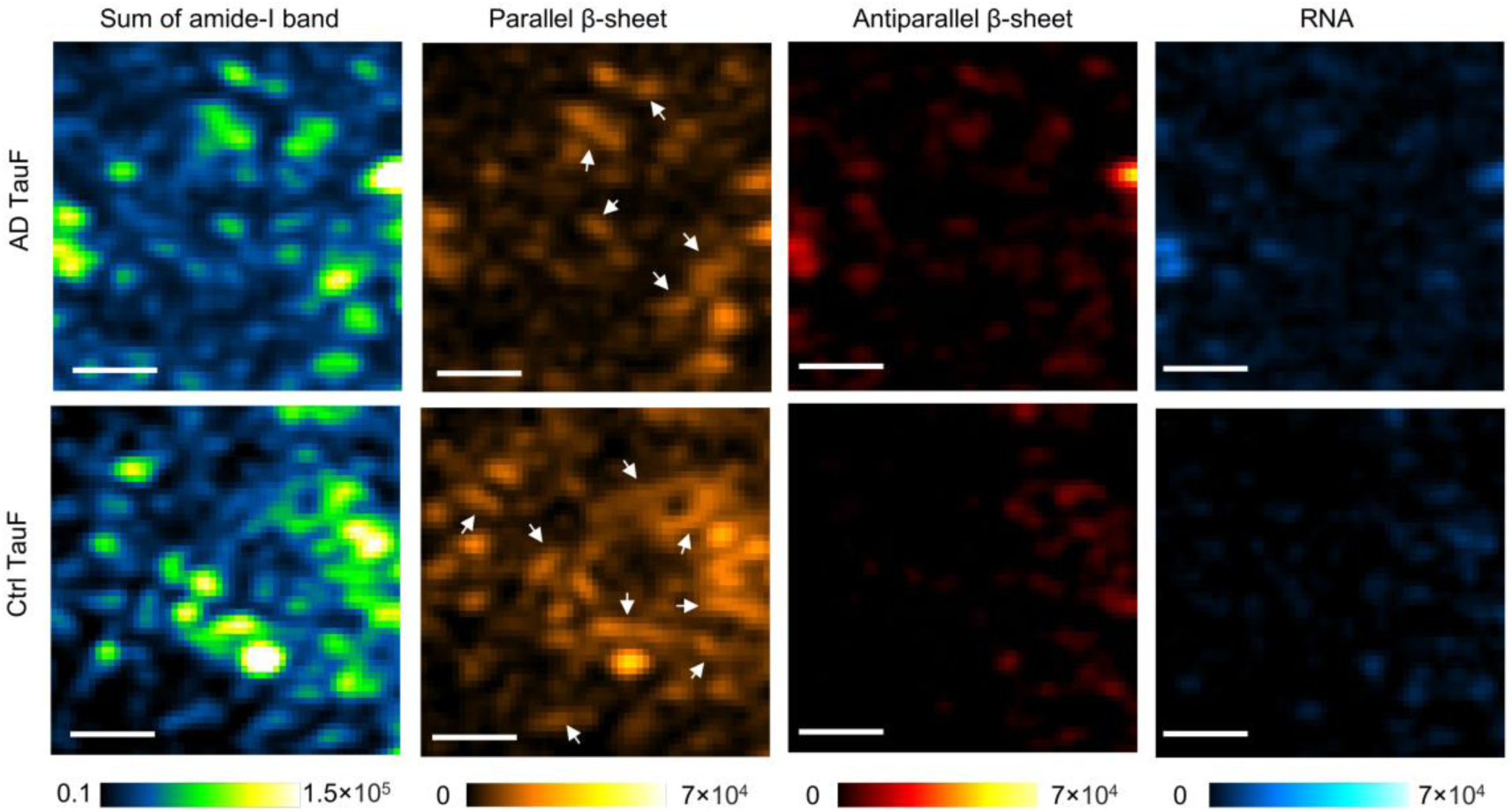
IR-AMES images of AD TauF and Ctrl TauF at the amide-I band. Sum of amide-I was obtained by integrating signals from 1,580–1,720 cm^−1^. Parallel β-sheet content was mapped using the 1,620–1,634 cm^−1^ band, antiparallel β-sheet using 1,680–1,694 cm^−1^, and RNA-associated signatures using the uracil C=O band using 1,694–1,710 cm^−1^. White arrows indicate representative elongated fibrillar structures. Scale bars: 1 µm. Additional morphological and spectroscopic characterization of diluted AD TauF samples is shown in **Supplementary** Fig. 24, confirming that AD TauF predominantly exists as short fibrillar assemblies with consistent spectral signatures.

**Extended Data Fig. 5.**
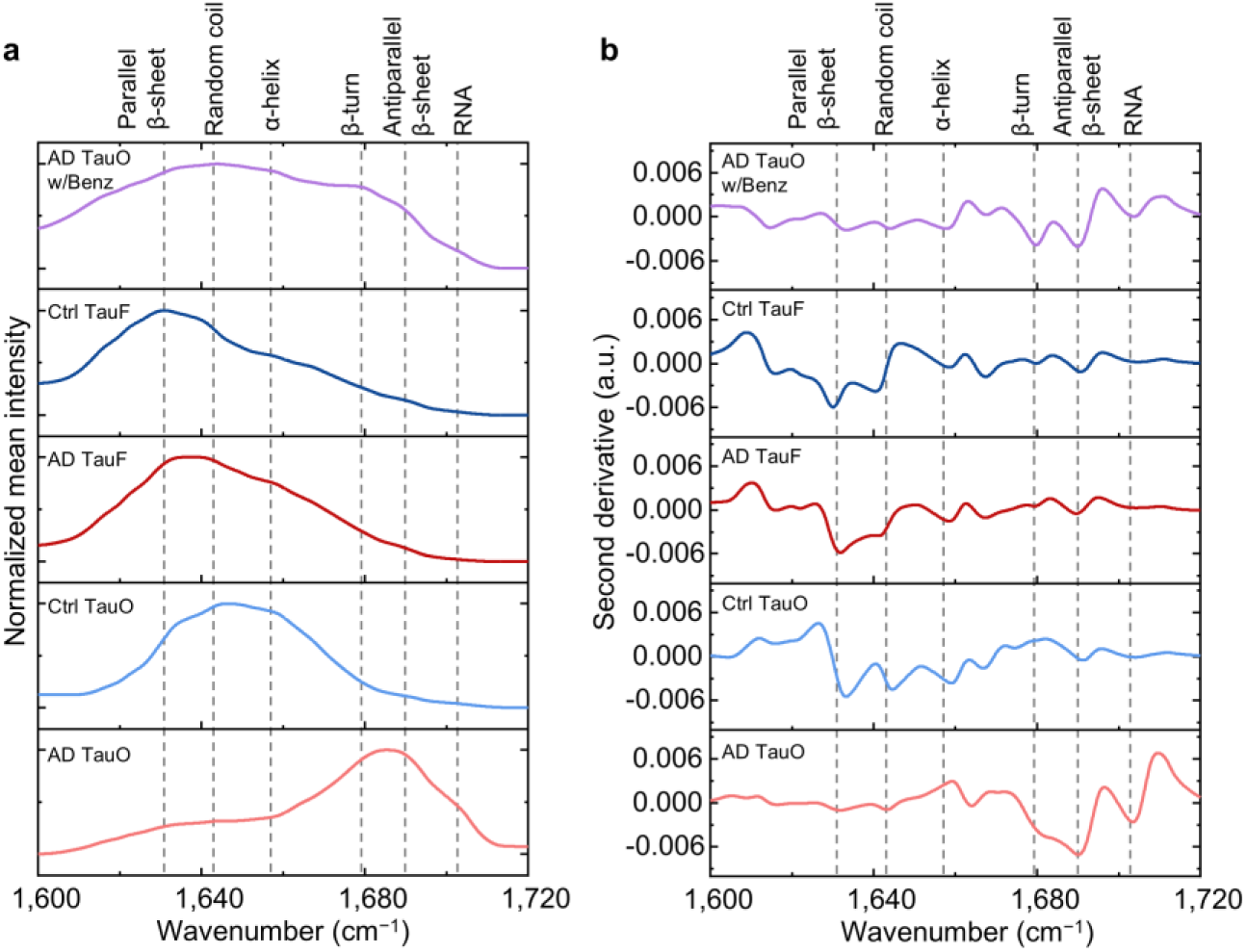
Amide-I spectral features and second-derivative analysis of human brain-derived tau assemblies. a, Mean amide-I spectra of AD TauO, Ctrl TauO, AD TauF, Ctrl TauF and benzonase-treated AD TauO. **b,** Corresponding second-derivative spectra, highlighting spectral minima associated with parallel β-sheet, random coil, α-helix, β-turn, antiparallel β-sheet and RNA-related vibrational features. Dashed lines indicate reference spectral positions used to guide component-center assignment for subsequent spectral decomposition.

**Extended Data Fig. 6.**
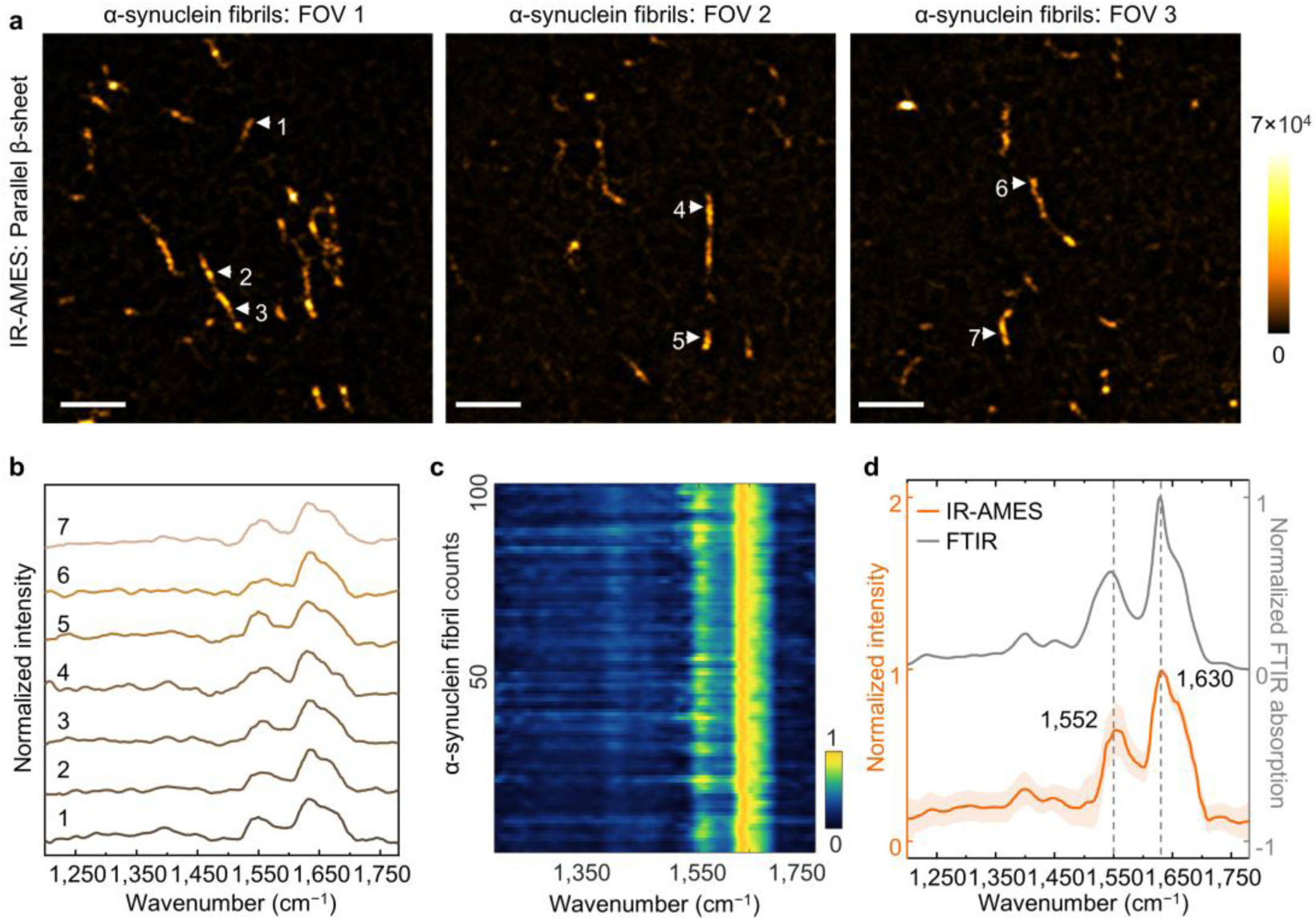
IR-AMES imaging and spectral characterization of in vitro–formed α- synuclein amyloid fibrils in Tris buffer. a, Representative IR-AMES images of α-synuclein fibrils acquired in the parallel β-sheet channel from three independent fields of view, showing elongated micrometre-scale fibrillar morphologies. Scale bars: 5 µm. b, Representative single-fibril IR-AMES spectra extracted from the positions indicated in a. c, Heatmap of normalized IR-AMES spectra collected from individual α-synuclein fibrils (*n* = 100). d, Comparison of the mean IR-AMES spectrum with the ensemble FTIR absorption spectrum, showing characteristic amyloid-associated spectral features, including the parallel β-sheet band near 1,630 cm^−1^. Solid lines: mean spectra, shaded regions: standard deviation.

