## Supporting Information for "Structural and compositional profiling of individual biomolecular assemblies by infrared absorbance-modulated evanescent scattering"

**Number of Pages:**

**Pages 2-11: Supplementary Notes 1-9**

**Pages 12-15: Supplementary Tables 1-4**

**Pages 16-49: Supplementary Figs. 1-34**

**Page 50: Supplementary References**

### Supplementary Note 1. Electromagnetic simulations supporting IR-AMES.

Electromagnetic simulations were performed to elucidate the physical mechanisms underlying the enhanced photothermal sensitivity of IR-AMES. Specifically, simulations were conducted to:

(i) Calculate mid-infrared field enhancement on the gold-coated substrate;
(ii) Compare visible probe-field distributions in coaxial and orthogonal-evanescent geometries;
(iii) Estimate the photothermal-induced scattering modulation.

#### 1.1 Mid-IR field enhancement

Mid-IR field distributions were simulated using Ansys Lumerical FDTD Solutions (v2024 R2.1). The model consisted of a 50-nm-thick gold film on a glass substrate, with air above the film. A monochromatic plane wave at 1,728 $cm^{-1}$, corresponding to the PMMA C=O vibrational band, was introduced using a total-field scattered-field source propagating along the $z$ axis toward the substrate. To study the dependence of the field distribution on illumination geometry, simulations were performed for incident angles $\theta = 0°$, 15°, 30°, and 45° within the $x$-$z$ plane. Polarization was controlled by the source polarization angle: 0° for $p$ polarization (electric field in the plane of incidence) and 90° for $s$ polarization (electric field perpendicular to the plane of incidence). The incident field amplitude was set as $|E_0|^2 = 1$. Perfectly matched layers were applied on all boundaries. The spatial distribution of the electric-field intensity was recorded using a frequency-domain 2D profile monitor, providing $x$-$z$ maps of the normalized field intensity $|E_{IR}|^2/|E_0|^2$ above the gold surface.

As shown in **Supplementary Fig. 1**, stronger interference-induced IR field enhancement is observed at 0°, 15°, and 30°, but the enhanced intensity is primarily distributed away from the interface (>1 μm) and therefore contributes minimally to excitation within the near-surface detection region (<100 nm). By contrast, 45° incidence ($p$ polarization) yields a modest (~2×) but spatially confined interfacial enhancement that is optimally matched to nanoscale measurements.

#### 1.2 Visible probe-field distribution

To compare conventional coaxial interferometric photothermal detection with the orthogonal-evanescent geometry used in IR-AMES, electromagnetic simulations were carried out using COMSOL Multiphysics 6.0 based on previous work[1]. A PMMA particle with diameter $d_{PMMA}$ of 500 nm was positioned at the interface of air and gold-coated glass. The contact region between the particle and substrate was modelled as a circular area with a diameter of $0.4d_{PMMA}$. The refractive index of PMMA and glass were set to 1.4998 and 1.5253, respectively. The gold film was modelled with a thickness of 50 nm, using the dielectric function reported by Olmon et al.[2] The probe field was defined as a 450 nm plane wave incident from the glass substrate. For the orthogonal evanescent configuration, the incidence angle was set to 42°, corresponding to total internal reflection (TIR), whereas normal incidence at 0° was used for the coaxial configuration. Periodic boundary conditions were applied to the lateral boundaries of the simulation domain to emulate an extended interface. The simulation domain was surrounded by perfect matched layer to reduce back-reflections.

As shown in **Supplementary Fig. 2**, TIR generates a laterally propagating evanescent field confined to the interface and decaying exponentially with distance from the surface. This surface confinement enhances the local probe intensity at the particle–substrate interface by approximately 6.5-fold relative to coaxial illumination, thereby increasing interferometric scattering contrast.

#### 1.3 Photothermal modulation

To compare the photothermal-induced modulation between coaxial and orthogonal detection geometries, electromagnetic simulations were carried out in COMSOL Multiphysics 6.0 using a Multiphysics model coupling heat transfer and electromagnetic wave. Transient temperature evolution was first calculated using the Heat Transfer in Solids physics module. The heat source was confined to the PMMA particle and defined by an absorption cross-section of $4.64 \times 10^{-10} \mathrm{cm}^2$, driven by a 1 μs square heating pulse with a peak power of 0.4 W. The resulting temperature distribution and temporal dynamics were were passed to the following electromagnetic simulation steps, accounting for thermally induced optical perturbations.

The scattering-field modulation induced by photothermal effects was modelled using the Electromagnetic Waves module, incorporating both thermo-optic refractive-index changes and thermal expansion of the particle. The power flow of the scattered field was integrated on a surface located 1.2 μm away from the particle, within an 80° collection cone, with a barrier blocking the reflection path to replicate the numerical aperture of the experimental objective used in experiments. The scattering power was obtained by integrating the time-averaged Poynting flux over the defined monitoring surface corresponding to specific collection solid angles. Photothermal modulation of scattered power was evaluated by perturbing (i) the refractive index ($n$) of PMMA using a thermo-optic coefficient $-1 \times 10^{-4} \mathrm{K}^{-1}$, and (ii) the particle size ($r$) using a thermal expansion coefficient $1 \times 10^{-4} \mathrm{K}^{-1}$ scaled by the simulated temperature rise.

In these simulations, identical mid-IR excitation and visible probe parameters were used for the coaxial and orthogonal geometries to enable a direct comparison. Coaxial detection results in partial cancellation between thermo-optic and thermal expansion contributions, yielding a modulation depth of ~2.6% (**Supplementary Fig. 3a**). This value is larger than the ~0.2% modulation reported previously for coaxial detection[3], likely due to differences in excitation conditions. Although the same mid-IR QCL was used, the repetition rate and average power delivered to the sample differed between implementations. In IR-AMES, the on-sample peak power was optimized and was approximately 13-fold higher than in the earlier reported coaxial configuration. This improvement produced a larger transient temperature rise and thus a stronger absolute photothermal response (Details in **Supplementary Note 3)**. Importantly, this increase in excitation strength does not eliminate the intrinsic cancellation mechanism of coaxial detection.

In contrast, IR-AMES employs orthogonal evanescent scattering detection, which blocks specular reflection and collects scattering over an asymmetric angular range (42°–83°), eliminating cancellation between the two photothermal components (**Supplementary Fig. 3b**). This orthogonal collection geometry increases the modulation depth to ~−16.5%. Importantly, the increased modulation depth does not arise from a reduction in the overall scattering signal, where the scattering intensity change is of comparable magnitude in both geometries. Thus, IR-AMES increased the modulation depth by more than two orders of magnitude compared with coaxial detection.

**Supplementary Note 2. Correction of effective diameter in the evanescent field.**

The evanescent field decreases exponentially at *z*-direction from the surface into the medium[4], the penetration depth ($l$) of the evanescent field can be calculated by

$$l = \frac{\lambda}{4\pi}(n_1^2 sin\theta - n_2^2)^{-1/2}$$

where $\lambda = 450$ nm is the probe light wavelength, $\theta$ is the incident angle, $n_1 = 1.5253$ and $n_2 = 1.00028$ are the refractive index of the glass substrate and air medium at 450 nm, respectively. $l = 176.6$ nm when $\theta$ is set as 42°.

The scattering of the evanescent field by a nanoparticle depends on the distance ($z$) from the surface[5]. For particles with different sizes, the effective scattering diameter $D_{eff}$ and volume $V_{eff}$ of the particle can be given by

$$V_{eff} = \frac{4\pi}{3}(\frac{D_{eff}}{2})^3 = \int_0^D \pi(Dz - z^2)e^{-\frac{z}{l}}dz$$

where $D$ is the diameter of the particle. Taking this z-distance dependence into account, the effective diameters of 95, 70, 50 nm PMMA nanoparticles used in **Fig. 1h** should be 86.98, 65.87, 47.45 nm, respectively.

**Supplementary Note 3. Simulation of the temperature rise.**

To quantitatively understand the photothermal heating process in IR-AMES, we performed time-dependent thermal simulations using COMSOL Multiphysics 6.0 based on previous work[3]. The thermal diffusion under mid-IR excitation was simulated via the heat-transfer-in-solids module by solving the heat diffusion equation:

$$C_p \rho \frac{\partial T}{\partial t} + \nabla \cdot (-k \nabla T) = Q(t)$$

where $Q(t)$ represents the volumetric heat source arising from infrared absorption, $T$ is the temperature, $t$ is the time, $C_p$ is the heat capacity, $\rho$ is the density, and $k$ is the thermal conductivity of the material in the system.

For simulations in air, a single PMMA bead (diameter 500 nm or 50 nm) was placed on a 50-nm-thick-gold-coated glass substrate. The IR excitation was set to the PMMA C=O resonance at 1,728 $cm^{-1}$. The IR illumination was modeled as an elliptical beam with a semi-major axis a = 25 µm and semi-minor axis b = 22.85 µm, as measured experimentally (**Supplementary Fig. 12**). The average IR power was set to 20 mW and calibrated by the absorption cross section of PMMA at the selected wavenumber. The pulse repetition rate was 100 kHz with a pulse width of 1 µs. To consider reflective enhancement from the gold film, an IR field enhancement factor of 2 was applied. The initial temperature and all thermal boundaries were set to 298 K. Heat convection was neglected. The heat source term was defined within the PMMA bead volume. The simulated temperature distributions and transient heating profiles are shown in **Supplementary Fig. 13**. The maximum temperature rise ($\Delta T$) was calculated to be ~36.7 K for a 500-nm PMMA bead and ~0.41 K for a 50-nm bead under these conditions.

For simulations in aqueous environments (**Extended Data Fig. 2a**), a 100-nm PMMA bead was placed on the same gold-coated substrate and covered by a 110-nm-thick water layer. Simulations were performed at both the PMMA C=O band (1,728 $cm^{-1}$) and the water O–H bending band (1,644 $cm^{-1}$). The average IR powers were set to 11.8 mW (500 ns) and 2.7 mW (80 ns) at 1,728 $cm^{-1}$, and 15.5 mW (500 ns) and 3.5 mW (80 ns) at 1,644 $cm^{-1}$. The simulated temperature profiles (**Extended Data Fig. 2a**) show that long IR pulses (500 ns) induce strong bulk water heating, which overwhelms the particle signal. In contrast, short IR pulses (80 ns) confine heating to the nanoparticle, strongly suppress water absorption.

**Supplementary Note 4. Experimental validation of temperature response.**

To experimentally validate the temperature response under mid-IR heating in IR-AMES, we performed fluorescence thermometry[6, 7] using 500-nm PMMA beads labeled with fluorescein isothiocyanate isomer I (PMMA-FITC). PMMA-FITC beads were synthesized by reacting amine-functionalized PMMA beads with FITC via its isothiocyanate group (**Supplementary Fig. 31**).

Temperature calibration was performed using widefield epi-fluorescence microscopy (IX71, Olympus, **Supplementary Fig. 14a**). PMMA-FITC beads were dispersed in a glass-bottom dish in air and placed on a temperature-controlled heating pad, while the sample temperature was monitored by a thermal camera (FLIR A325sc). Fluorescence images were acquired using an air objective (Plan Fluor 20×/0.50, AmScope) and a scientific CMOS camera (ORCA-Flash4.0 V3, Hamamatsu; exposure time 5 s). The normalized fluorescence intensity decreases linearly with temperature increase, with a calibration slope of –0.23%/ K ($R^2$ = 0.9995).

To quantify the photothermal temperature rise under IR-AMES, the 500-nm PMMA-FITC beads were dispersed on a gold-coated glass slide and imaged at the single-particle level in the system. Fluorescence images were recorded with an exposure time of 45 ms (gain 30, frame rate 12.5 fps). The visible probe was the same light used in IR-AMES measurements, with a neutral-density filter applied to attenuate fluorescence excitation intensity and minimize photobleaching. Fluorescence detection was performed using a 490-nm long-pass dichroic mirror (DMLP490R, Thorlabs), a 500-nm short-pass excitation filter, and a 500-nm long-pass emission filter. The mid-IR pump laser operated at 100 kHz with a pulse width of 1 µs and was tuned to the PMMA C=O resonance (1,728 $cm^{-1}$). Under IR-on conditions, the fluorescence intensity of individual PMMA-FITC beads decreased by ~8.5% relative to IR-off frames (**Supplementary Fig. 14b**). Thus, the observed fluorescence modulation corresponds to a local temperature rise of approximately 37 K for 500-nm PMMA beads in IR-AMES detection, which matches the simulation well (**Supplementary Fig. 13a**). This also indicates that the orthogonal-scattering-based photothermal geometry (**Extended Data Fig. 1b**) provides ~2.5-fold larger modulation depth than fluoresce-detected photothermal redout.

**Supplementary Note 5. Benchmarking IR-AMES against representative O-PTIR modalities.**

To benchmark the performance of IR-AMES, we compared its photothermal modulation contrast with representative widefield and point-scanning optical photothermal infrared (O-PTIR) implementations reported using 500-nm PMMA particles. Since different O-PTIR detection schemes rely on distinct substrates (e.g., $CaF_2$, silicon, glass, or gold substrates), illumination geometries and optical configurations, measurements of the identical physical sample across platforms are generally not feasible. We therefore compare results obtained using the same type and size of reference particle under the reported operating conditions of each platform.

In pump–probe photothermal microscopy, infrared-induced absorption is converted into a modulated optical signal. As a result, the modulation depth ($\Delta I/I$) serves as a natural metric to evaluate sensitivity. The comparison is summarized below and in **Supplementary Fig. 8**.

1. **Widefield dark-field scattering detection.** Conventional coaxial widefield scattering-based mid-infrared photothermal detection[3] produced a modulation depth of approximately 0.2% from a single 500-nm PMMA particle using QCL excitation. Under comparable QCL excitation, IR-AMES achieves a modulation depth of approximately 22.7%, corresponding to more than two orders of magnitude enhancement in relative modulation contrast.
2. **Widefield interferometric detection.** Widefield interferometric defocus-enhanced mid-infrared photothermal microscopy[8] has achieved high sensitivity through interferometric scattering detection combined with high-pulse-energy OPO excitation. For single 500-nm PMMA particles, a modulation depth of approximately 7.6% has been reported. IR-AMES reaches approximately 22.7% modulation while using approximately tenfold lower IR pulse energy (**Supplementary Table 1**), illustrating the strong signal enhancement provided by the orthogonal evanescent-scattering geometry.
3. **Fluorescence-detected photothermal imaging.** Fluorescence-detected mid-infrared photothermal microscopy[9, 10] can generate strong photothermal contrast through temperature-dependent fluorescence modulation. For a direct comparison on the IR-AMES platform, we performed total internal reflection fluorescence (TIRF)-based photothermal measurements of 500-nm PMMA-FITC particles under comparable excitation conditions. The fluorescence intensity modulation was approximately 8.5% (**Supplementary Fig. 14b**), compared with approximately 22.7% scattering modulation measured by IR-AMES. We note that fluorescence-based approaches may achieve higher sensitivity depending on dye properties[11]. In contrast, IR-AMES operates in a label-free manner while maintaining strong modulation depth.
4. **Point-scanning O-PTIR detection.** Representative point-scanning O-PTIR measurements of single 500-nm PMMA particles have reported modulation depths of approximately 1–3%[12, 13]. Although point-scanning and widefield methods differ substantially in illumination, detection and acquisition strategy, these reported values provide an additional reference for the modulation contrast achieved by IR-AMES.

Taken together, these comparisons indicate that IR-AMES provides high photothermal modulation contrast for label-free nanoparticle detection while retaining widefield imaging capability.

**Supplementary Note 6. AlphaFold3 structure predictions of recombinant human 2N4R tau monomer and oligomers.**

Structural models of recombinant human 2N4R tau monomers (rTauM) and oligomers (rTauO) were generated using the AlphaFold Server (AlphaFold3). For rTauM, a single protein entity was used, whereas rTauO, two or three identical protein entities were specified to model dimeric and trimeric assemblies, respectively. All predictions were performed using the full-length human tau sequence (2N4R isoform), listed below:

MAEPRQEFEVMEDHAGTYGLGDRKDQGGYTMHQDQEGDTDAGLKESPLQTPTEDGSEEPGSE
TSDAKSTPTAEDVTAPLVDEGAPGKQAAAQPHTEIPEGTTAEEAGIGDTPSLEDEAAGHVTQAR
MVSKSKDGTGSDDKKAKGADGKTKIATPRGAAPPGQKGQANATRIPAKTPPAPKTPPSSGEPPK
SGDRSGYSSPGSPGTPGSRSRTPSLPTPPTREPKKVAVVRTPPKSPSSAKSRLQTAPVPMPDLKNV
KSKIGSTENLKHQPGGGKVQIINKKLDLSNVQSKCGSKDNIKHVPGGGSVQIVYKPVDLSKVTS
KCGSLGNIHHKPGGGQVEVKSEKLDFKDRVQSKIGSLDNITHVPGGGNKKIETHKLTFRENAKA
KTDHGAEIVYKSPVVSGDTSPRHLSNVSSTGSIDMVDSPQLATLADEVSASLAKQGL.

Tau is a highly intrinsically disordered protein. Consequently, AlphaFold3 predictions are expected to display substantial conformational variability and limited confidence compared with folded proteins. The predicted structures are therefore not intended to represent unique or definitive conformations, but instead serve as qualitative structural references for conformational heterogeneity and intermolecular organization.

To sample conformational variability, multiple independent AlphaFold3 runs were performed using distinct automatically generated random seeds, listed below:

rTauM seeds: 425710735, 1040918707, 275144931, 659470081, 1739733549, 2034550252, 629307008, 1881351104, 1768348907, 307590870, 2131491728, 1750398382, 382668278, 022368216, 1021227079, 136257710, 220629208, 411064369, 455245725, 2065531763.

rTauO (dimer) seeds: 2114177173, 406531529, 1486848431, 1632499544, 251760084, 2081774202, 841200492, 987097790, 1833961065, 143373696.

rTauO (trimer) seeds: 98037787, 1371087947, 1796823790, 697333908, 1184016740, 30008249, 510686364, 1824008212, 1676206883, 1784508607.

For tau monomers, repeated predictions revealed heterogeneous conformations consistent with their intrinsically disordered nature, dominated by random-coil-like features with short α-helix structures (**Supplementary Fig. 18a**). In contrast, predicted dimeric and trimeric assemblies frequently showed β-sheet-rich structures within disordered conformations (**Supplementary Fig. 18b**), leading to coexistence of β-sheet-rich, partially folded, and highly disordered conformational states within early tau oligomers.

**Supplementary Note 7. Sample preparations for tau and nanodisc samples.**

Tau oligomers and fibrils are conformationally sensitive. To minimize perturbation of native structures, tau samples were immobilized on substrates via nonspecific adsorption without further surface modification. Incubation times of tau assemblies, nanodiscs, and tau–nanodisc complexes ranged from 15 min to 2 h depending on sample concentration and adsorption kinetics and were adjusted until sufficient individual particles were stably adsorbed on the surface. Validation is provided in **Supplementary Fig. 30** to exclude motion-induced artifacts in the solution condition.

Recombinant human 2N4R tau monomer (rTauM, 46 kDa, 2.2 mg $mL^{-1}$ in PBS, pH 7.4) was diluted to 8 µM in a 160-µL reaction containing 10 mM HEPES, 100 mM NaCl, 5 mM DTT and 0.1 mM EDTA. Tau oligomerization[14] was initiated by adding freshly prepared arachidonic acid (AA) to a final concentration of 300 µM. The solution was gently mixed and briefly centrifuged (3–5 s). The resulting tau oligomers were protected from light, and diluted to 10 nM for immediate IR-AMES measurements. For imaging, diluted tau oligomers were deposited onto cleaned gold-coated glass slides, gently rinsed with buffer to remove unbound species, and sealed in a thin aqueous layer using a $CaF_2$ coverslip, as described in the **Methods**.

Human-derived tau samples (40 ng/µL) were prepared as pooled mixtures from postmortem brains: Ctrl TauO, AD TauO, Ctrl TauF and AD TauF. For IR-AMES measurements, tau samples were diluted to 4 ng/µL, and deposited on cleaned substrates, gently rinsed with buffer to remove unbound material, and sealed in a thin aqueous layer with a $CaF_2$ coverslip.

To assess RNA contributions, AD TauO were first adsorbed onto cleaned gold-coated glass slides by incubating the substrate with 2 µL of AD TauO solution (4 ng/µL) at 4 °C. After gentle rinsing with buffer, the surface was incubated for 30 min with enzyme buffer consisting of 20 mM Tris-HCl, 0.1 U/µL Benzonase, and 2 mM $MgCl_2$. Samples were rinsed 3-5 times with buffer, then covered with a thin layer of buffer and sealed with a $CaF_2$ coverslip for IR-AMES measurements.

Nanodiscs were prepared in 25 mM Tris (pH 8.0) and 100 mM NaCl, and used at working concentrations of 150 nM. For tau–nanodisc interaction studies, tau oligomers or fibrils (50 nM) were mixed with nanodiscs (150 nM) in Tris/NaCl buffer, with the addition of $ZnCl_2$ to a final concentration of 50 µM to enhance tau–membrane association. The total reaction volume was 20 µL. Samples were gently mixed and incubated for 30 min at room temperature prior to imaging. Nanodiscs alone and tau–nanodisc mixtures were deposited onto cleaned gold-coated glass slides and allowed to adsorb naturally at room temperature. After incubation, the substrates were gently rinsed with Tris/NaCl buffer to remove unbound particles. Samples were rinsed 3–5 times with buffer, then covered with a thin layer of buffer and sealed with a $CaF_2$ coverslip for IR-AMES measurements.

**Supplementary Note 8. IR-AMES image and spectral processing workflow.**

IR-AMES images are generated by subtracting the scattering image at IR-off from IR-on frames at each wavenumber, followed by background correction and spatial bandpass filtering (ImageJ, 2–2000 pixels; pixel size 75 nm) to suppress pixel-level noise and large-scale background while preserving diffraction-limited particle features (~200 nm) (**Supplementary Fig. 6**). The resulting images and spectra were processed as described below.

Individual nanoparticles were automatically identified using TrackMate[15] plugin in ImageJ with a fixed spot diameter of 3–4 pixels (~225–300 nm, camera pixel size 75 nm) (**Supplementary Fig. 32**), matching the diffraction-limited spot size of the imaging system (**Fig. 1i**). For particles larger than the diffraction limit, such as 500-nm PMMA beads, their physical size was used for particle selection. For each identified particle, IR-AMES spectra were extracted by averaging the pixel intensities within the selected region at each wavenumber.

Extracted spectra were baseline-corrected, normalized by the measured IR power at each wavenumber, and lightly smoothed (7–11 points) to suppress noise without distorting spectral features. Water absorption along the IR beam path can induce sharp power dips at specific wavenumbers, resulting in abnormally low IR power (**Supplementary Fig. 33a**). To prevent artifacts arising from normalization by these low-power points, wavenumbers strongly affected by water absorption were identified from the measured IR power spectrum and excluded from further analysis (**Supplementary Fig. 33b**). The remaining spectra were then normalized by the corresponding IR power to correct for wavelength-dependent power variation (**Supplementary Fig. 33c, d**).

Owing to the coherent interferometric scattering from the nanoscale roughness of the substrate[5, 16], IR-AMES images can contain speckle-like or line-like background features, which may become more apparent in chemical-band-integrated images. For displaying a single chemical component, representative single-wavenumber images at the corresponding peak position were used when appropriate, whereas quantitative spectral analysis was performed on the full hyperspectral stack. Automatically identified spots were validated by their target-material vibrational signatures after spectral extraction and preprocessing. For example, in IR-AMES imaging of 95-nm PMMA nanoparticles, identified spots showed the expected C=O resonance near 1,730 cm$^{-1}$, whereas representative line-like or background features lacked this PMMA-specific signature (**Supplementary Fig. 34**). Therefore, objects were retained for single-particle spectral analysis only when they exhibited diffraction-limited morphology and the expected target-material vibrational features.

Spectral components were quantified by constrained multi-component fitting in MATLAB (**Supplementary Fig. 20**). For each sample group, single-particle spectra were averaged and smoothed using a Savitzky–Golay filter. The second derivative of the averaged spectrum was used to identify the major component bands, and the resulting band positions were used to constrain the fitting of individual single-particle spectra. Peak amplitudes were fitted as non-negative parameters, whereas peak centers were allowed only limited bounded shifts from the group-defined positions. Component areas were obtained by integration and used to calculate the relative contribution of each spectral component for every particle.

**Supplementary Note 9. Neuronal toxicity assay.**

**LDH assay.** 50 µL supernatant was collected as designed time point into a 96-well plate for lactate dehydrogenase (LDH) release assay as per manufacture's protocol. Briefly, 50 µL of the CytoTox 96® Reagent was added to each sample aliquot. The plate was covered with foil to protect it from light and incubated for 30 min at room temperature on shaker. 50 µL of Stop Solution was added to each well of the 96-well plate and the absorbance recorded at 490 nm with the plate reader. Each experiment was repeated at least three times with triplicate wells each time.

**Immunocytochemistry (ICC) staining of fixed iPSC-derived neurons**. iNeurons on a 24-well cover slips were fixed with 0.5 mL 4% PFA/PBS for 15 min. The cells were washed three times in PBS, 5 min each wash. The cells were permeabilized in .0.5 mL PBS/0.1% Triton X-100 (PBST) for 15–30 min. Blocking was done in 0.5mL of 5% BSA with 5% donkey Serum in PBST for 1 h. Then the cells were incubated in primary antibodies diluted in 5% BSA/PBST at 4 °C overnight followed by being washed 3 times in PBS-T, 10 min each, on the second day. The samples were incubated in 2° antibody diluted in 5% BSA/PBST, 2 h at RT. After incubation with the 2° antibody, the samples were incubated in DAPI diluted 1:10,000 in PBST (5 mg/mL stock solution) for 5 min after first wash. Then the samples were washed with 2 times with PBST, and then once in PBS, 10 min each, after which the samples were mounted onto coverslips using Prolong Gold Antifade mounting media. The primary antibodies used in this study for ICC are as follows: CP-13 (mouse, provided by The Feinstein Institutes for Medical Research, 1:300 dilution), Cleaved Caspase 3 (rabbit, Cell Signaling Technology, cat# 9661, dilution 1:500), Tuj1/Anti-Beta-Tubulin 3 Antibody (chicken, Aves Labs, cat# TUJ, dilution 1:500). All the 2° antibodies were purchased from Thermo Fisher Scientific made in donkey and used for 1:800 dilution in staining. Images were captured by Leica Stellaris 5 confocal microscope.

After 24 h treatment of iPSC-derived neurons ($n$ = 6), tau oligomers induced greater cytotoxicity compared with fibrils, as reflected by increased LDH release in oligomer-treated groups. Consistently, immunofluorescence analysis revealed elevated cleaved caspase-3 signals following oligomer exposure, indicating enhanced apoptotic activation. Within each tau conformational class, AD-derived tau produced stronger effects than age-matched control-derived tau. Oligomer treatment was also associated with increased CP13 immunoreactivity and reduced TUJ1-positive neurite area, consistent with increased tau phosphorylation and neurite disruption. These measurements collectively indicate stronger neurotoxic responses associated with AD tau oligomers relative to other species under the experimental conditions used in this study.

**Supplementary Table 1. Performance of IR-AMES and widefield interferometric defocus-enhanced mid-IR photothermal (WIDE-MIP)[8] imaging.** Comparison is performed using 500-nm PMMA beads in air at the C=O resonance 1,728 $cm^{-1}$. Signal-to-noise ratio was calculated as the mean intensity of single particle divided by the standard deviation of intensity in a background region.

| **System** | IR-AMES | WIDE-MIP |
|---|---|---|
| **Mid-IR source** | QCL laser<br>(MIRcat 2400, Daylight Solutions) | OPO laser<br>(Firefly-LW, M Squared Lasers) |
| **Pulse energy** | ~0.2 μJ | ~ 2.5 μJ |
| **Camera speed** | 200 fps | 1250 fps |
| **Chemical imaging speed** | 100 Hz, single frame (cold−hot) | 1.56 Hz, 400 frames (cold−hot) averaged |
| **Modulation depth** | ~22.7% | ~7.6% |
| **Signal-to-noise ratio** | 571 | 268 |

**Supplementary Table 2. Peak assignments in IR-AMES spectra.** The protein amide-I band arises mainly from C=O stretching vibrations, with the band position determined by the backbone conformation and the C=O···H–N hydrogen-bonding pattern[17]. In contrast, the amide-II band (1,480–1,580 $cm^{-1}$), which originates from C–N stretching and N–H bending vibrations, lacks structural specificity for secondary structure analysis. Lorentzian deconvolution was performed within the amide-I region to quantify protein secondary-structure components, RNA-associated carbonyl vibrations and lipid C=O stretching modes, enabling analysis of protein conformation and molecular composition at the single-particle level.

| **Fitting components** | **Peak range ($cm^{-1}$)** |
|---|---|
| low-frequency β-sheet (mainly parallel) | 1,615–1,634 |
| Random coil | 1,638–1,648 |
| α-helix | 1,650–1,660 |
| β-turn | 1,664–1,680 |
| high-frequency β-sheet (mainly anti-parallel) | 1,680–1,694 |
| RNA[18] (C=O stretching in Uracil) | 1,696–1,710 |
| Lipid[19] (C=O stretching) | 1,732–1,748 |

**Supplementary Table 3. Demographic, genetic and neuropathological characteristics of human brain donors.**

| Primary Neuropathologic Diagnosis | Braak Stage | Age at Onset | Age at Death/Bx | Duration (years) | APOE | Race | Sex |
|---|---|---|---|---|---|---|---|
| Alzheimer's disease (AD) | VI | 49 | 59 | 10 | E3/3 | w | m |
| AD | VI | 59 | 69 | 10 | E3/4 | w | m |
| AD | VI | 53 | 71 | 18 | E3/4 | w | m |
| AD | VI | 65 | 80 | 15 | E3/4 | b | f |
| AD | V | 77 | 83 | 6 | NA | w | f |
| AD | V | 86 | 92 | 6 | E3/4 | w | f |
| AD | V | 82 | 92 | 10 | E3/3 | w | m |
| AD | VI | 80 | 93 | 13 | E3/4 | w | f |
| Normal Control (Ctrl) | I | NA | 59 | NA | E2/3 | b | m |
| Ctrl | I | NA | 70 | NA | E3/3 | b | m |
| Ctrl | I | NA | 72 | NA | E3/3 | w | m |
| Ctrl | II | NA | 78 | NA | E3/3 | w | f |
| Ctrl | IV | NA | 84 | NA | E2/3 | b | f |
| Ctrl | III | NA | 91 | NA | E3/3 | w | f |
| Ctrl | III | NA | 92 | NA | E3/3 | w | f |
| Ctrl | II | NA | 94 | NA | E3/3 | w | m |

**Supplementary Table 4. Average probe power density and camera exposure time used for different measurements.** The average probe light power densities used in this work are comparable to those commonly employed in single-protein imaging studies[5, 20].

| Samples | Probe power density ($kW/cm^2$) | Camera exposure time (ms) |
|---|---|---|
| 500-nm PMMA particle | 0.1 | 4.5 |
| 95-nm PMMA particle | 0.15 | 9.5 |
| 70-nm PMMA particle | 0.18 | 14.5 |
| 50-nm PMMA particle | 0.25 | 14.5 |
| 38-nm PMMA particle | 0.25 | 14.5 |
| IgM (air) | 6 | 14.5 |
| Proteins and protein–nanodisc complexes in buffer (IgM, rTauM, rTauO, human TauO, TauF, α-synuclein amyloid fibrils) | 6 | 14.5 |

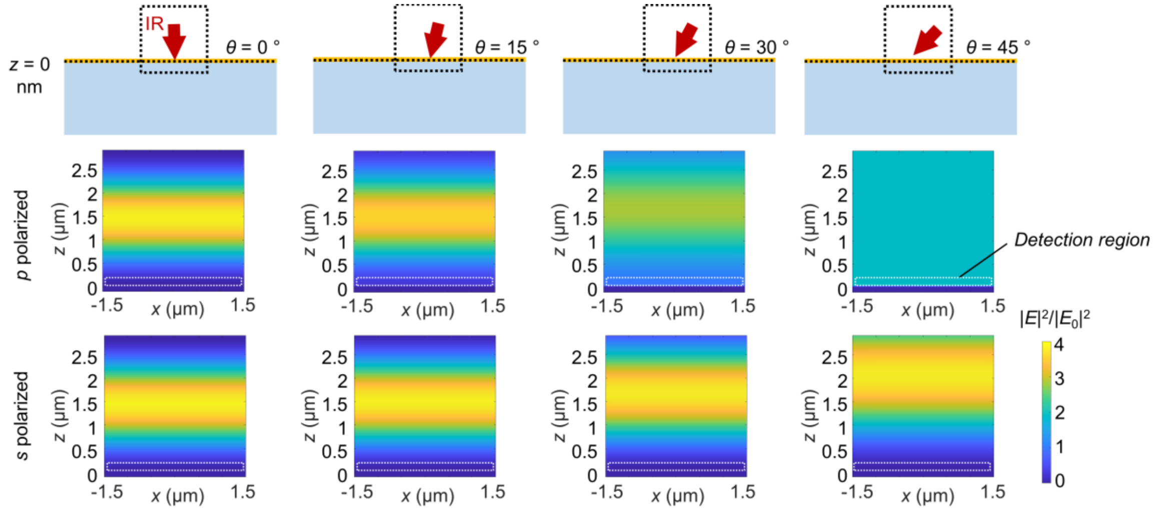


**Supplementary Fig. 1 Simulated IR intensity distributions at the air-gold interface under different incident angles ($\theta$).** The IR wavenumber corresponds to the C=O vibrational band (1,728 $cm^{-1}$). Images show the normalized IR intensity $|E|^2$ in the *xz* plane for both *p*- and *s*-polarized excitations. The incident IR intensity: $|E_0|^2 = 1$. The white dashed line indicates the detection region near the surface, showing that *p*-polarized excitation at $\theta = 45°$ produces the strongest IR field enhancement in the detection region.

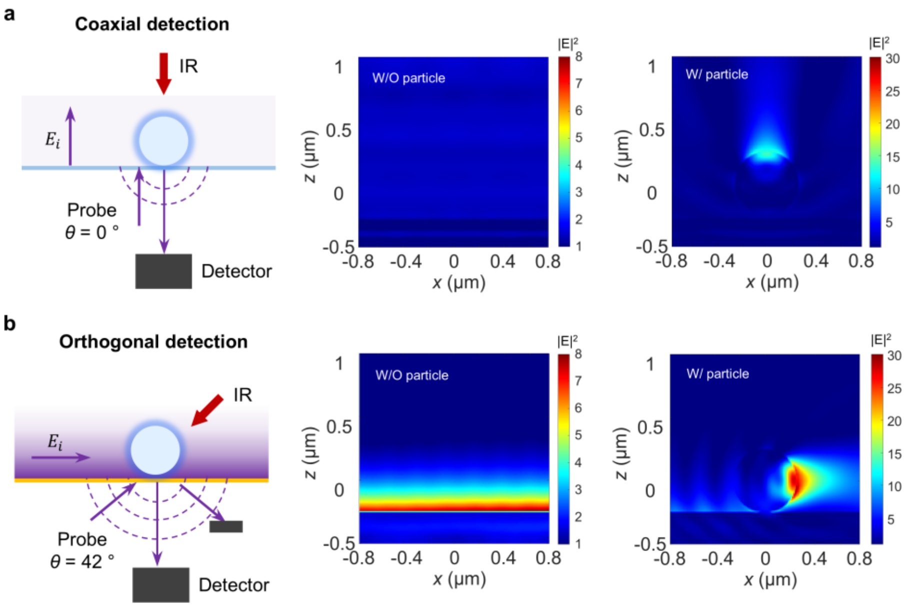


**Supplementary Fig. 2 Comparison of coaxial versus orthogonal detection schemes using simulated probe field distributions. a–b,** Detection geometries and simulated probe-field intensity distributions with (W/) and without (W/O) a 500-nm PMMA particle at the substrate–air interface for conventional detection of interferometric scattering (**a**) and orthogonal detection of TIR-illuminated evanescent scattering (**b**). In the orthogonal detection configuration, the evanescent field propagates laterally along the interface and decays exponentially with distance from the surface. This surface confinement increases the local probe intensity by ~6.5-fold. The incident light intensity: $|E_0|^2 = 1$.

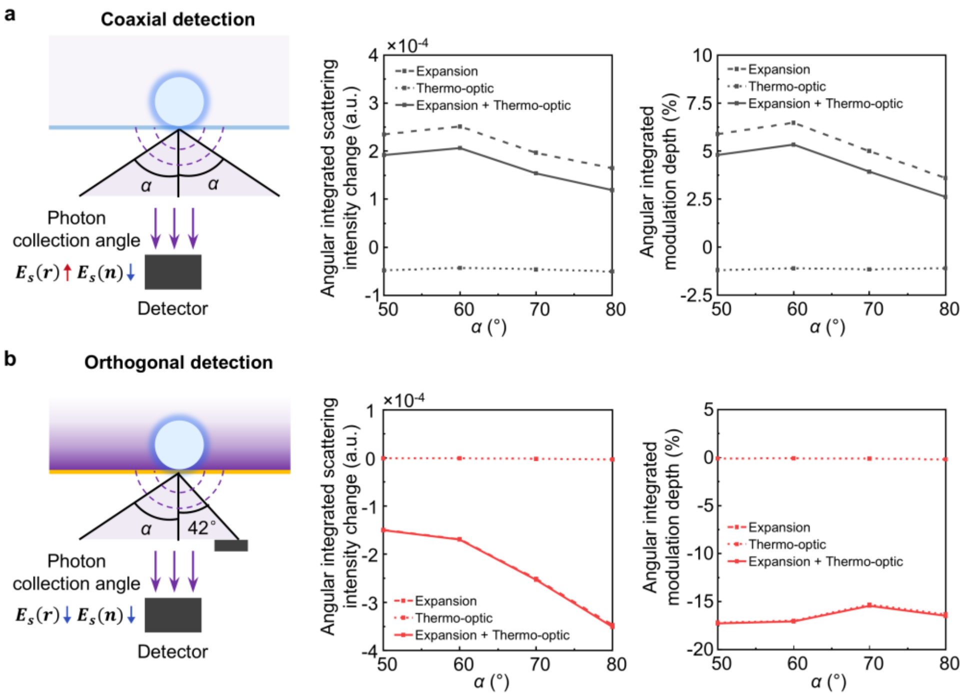


**Supplementary Fig. 3 Numerical simulations of photothermal induced scattering modulation of coaxial versus orthogonal-evanescent detection. a–b,** Detection geometries (left) and calculated angular integrated scattering intensity changes (middle) and modulation depths (right) of a 500-nm PMMA particle at the substrate–air interface for conventional coaxial interferometric photothermal detection (**a**) and orthogonal-evanescent photothermal detection (**b**). For coaxial detection with normal (0°) incidence, photon collection angle is $\alpha$ relative to the optical axis. For orthogonal detection with 42° TIR-illumination in air, reflected light is blocked to isolate the evanescent scattering signals, resulting in asymmetric collection angles, with 83° on the incident side, corresponding to the aperture of the oil-immersion objective, and 40° on the reflection side. The results indicate the partial cancellation between thermal expansion ($r$) and thermo-optic ($n$) effects in the coaxial geometry, and stronger modulation in the orthogonal detected TIR-illuminated evanescent scattering configuration.

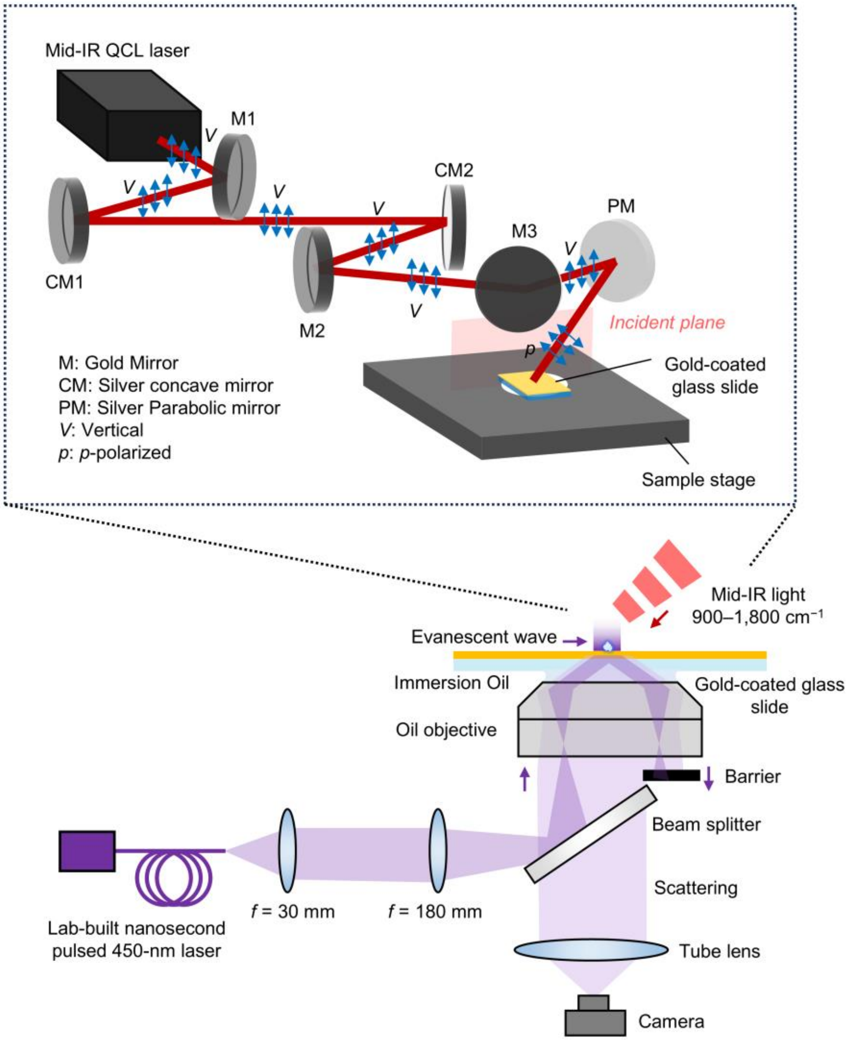


**Supplementary Fig. 4 IR-AMES setup.** The IR-AMES platform was constructed on an inverted microscope and integrated a TIR visible probe (bottom) with widefield mid-IR photothermal excitation (top). The visible probe was generated by a lab-built nanosecond pulsed 450-nm laser and delivered through a fiber. The mid-IR pump was provided by a tunable pulsed quantum cascade laser (QCL) covering 900–1,800 cm$^{-1}$. The IR beam was focused onto the gold-coated substrate using parabolic mirrors at oblique incidence. Additional details are provided in the **Methods**.

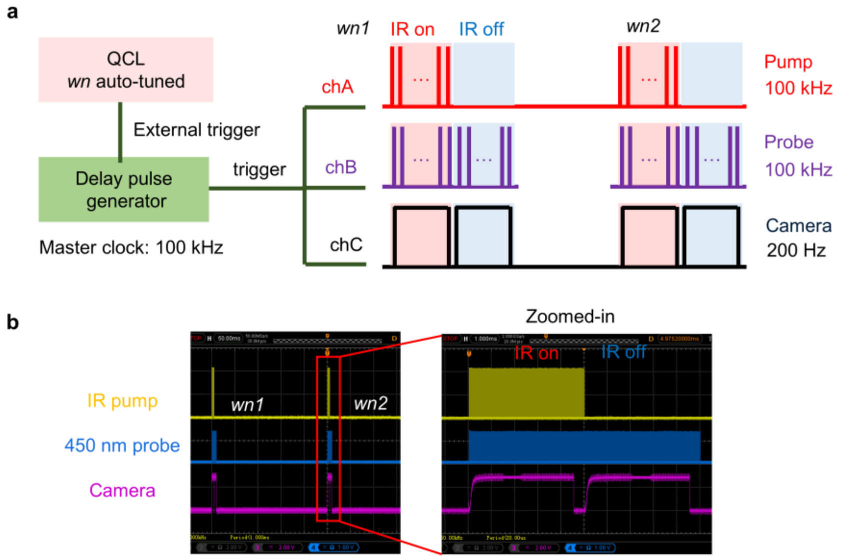


**Supplementary Fig. 5 IR-AMES signal acquisition and synchronization. a,** Schematic of the synchronization workflow. *wn*: IR wavenumber. During QCL auto-step scanning, the start of each wavenumber step sends an external trigger to a delay pulse generator. The generator, running with a 100-kHz master clock, synchronously triggers the mid-IR pump laser (100 kHz), the 450-nm probe laser (100 kHz), and the camera (200 Hz) to synchronize the IR pump pulses, visible probe pulses, and camera exposure at each wavenumber. This sequence repeats as the QCL scans through all programmed wavenumbers. The internal trigger step time determines the IR firing window per frame and is set to match the camera exposure. The camera is triggered to acquire both one IR-on (hot) frame and one IR-off (cold) frame at each wavenumber, and the IR-AMES photothermal contrast is obtained by subtracting cold from hot frames. **b,** Oscilloscope traces showing synchronized IR pump pulses (yellow), visible probe pulses (blue), and camera exposure pulses (purple) during wavenumber scanning.

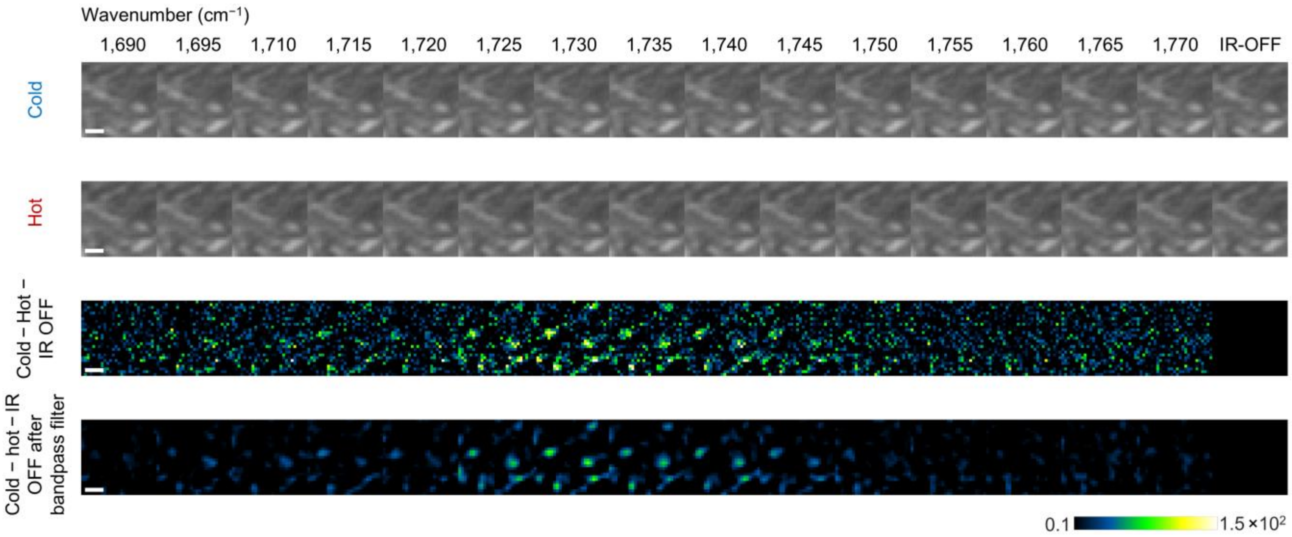


**Supplementary Fig. 6 IR-AMES image-processing workflow.** IR-encoded scattering images of 50-nm PMMA particles are generated by subtracting IR-off (cold) from IR-on (hot) frames during wavenumber scanning, followed by background correction and spatial bandpass filtering to suppress noise. The processed images are used to produce hyperspectral datasets. Scale bars: 500 nm.

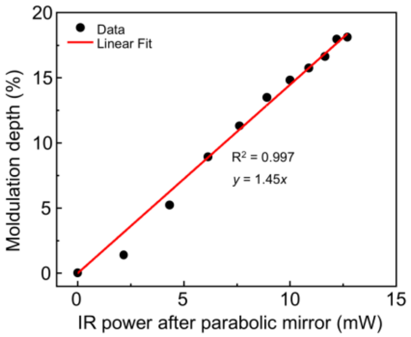


**Supplementary Fig. 7 Dependence of scattering modulation on mid-IR power.** The linear relationship between scattering change and IR absorption indicates minimal-system-induced nonlinear effects in the measurement.

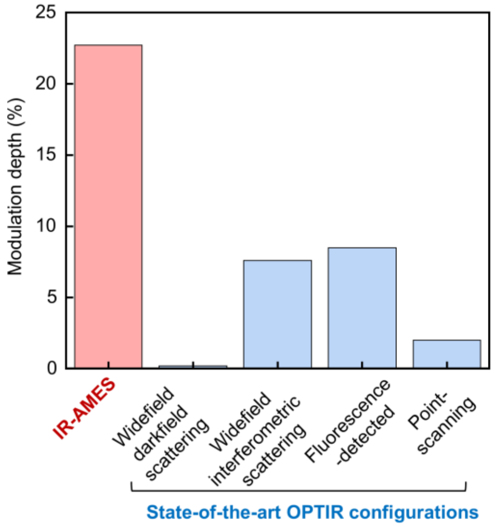


**Supplementary Fig. 8 Comparison of IR-AMES with representative OPTIR implementations.** Photothermal modulation depths measured from single 500-nm PMMA particles across representative OPTIR configurations. A detailed comparison is provided in **Supplementary Note 5**.

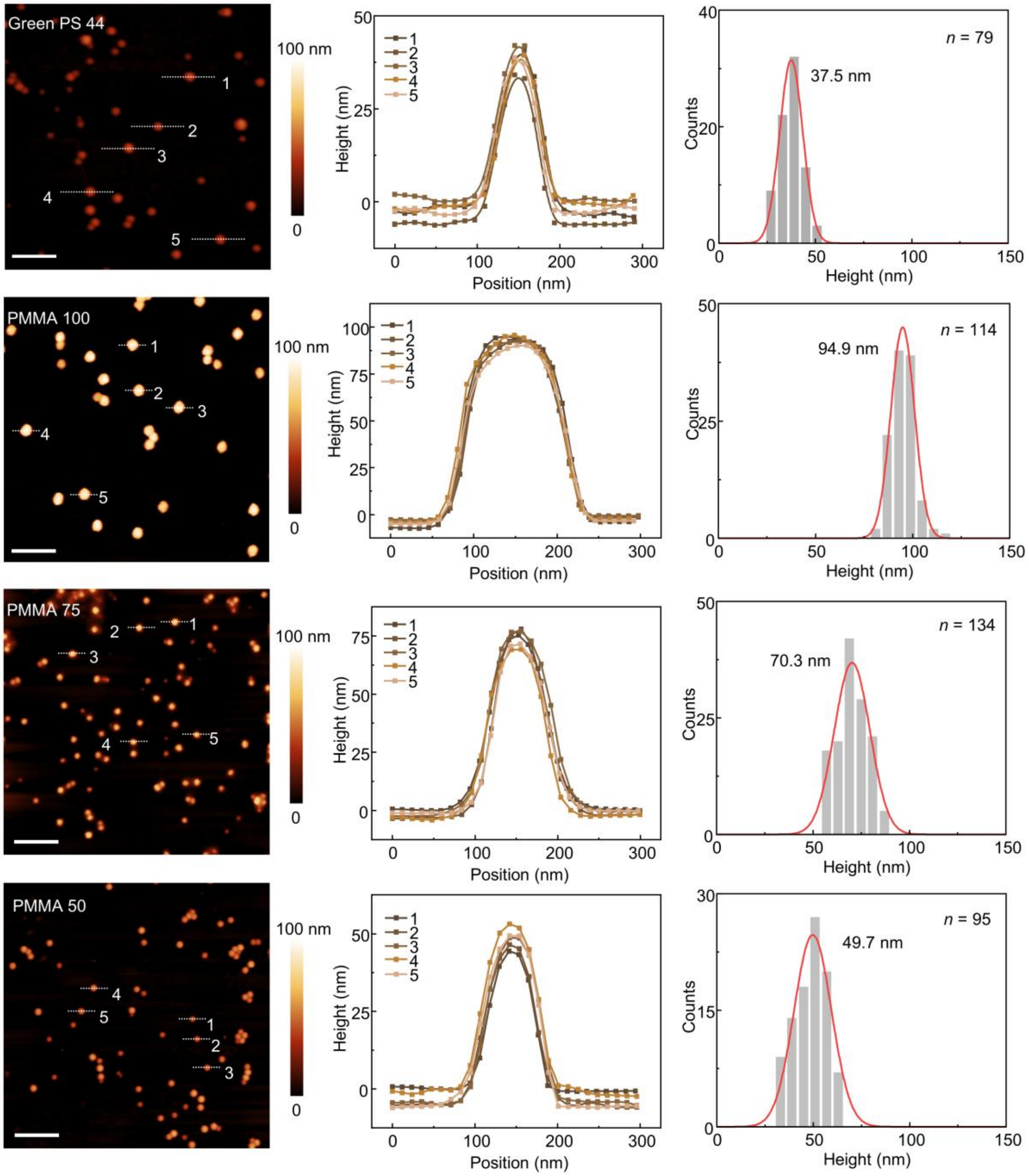


**Supplementary Fig. 9 Morphological characterization of standard nanoparticles by AFM.** AFM images, representative line profiles and height distributions of fluorescent PS and PMMA nanoparticles. The particles were well dispersed and exhibited bead-like morphologies. Five representative height profiles are shown for each group. Gaussian fitting of the height distributions yielded centers of 37.5, 94.9, 70.3, and 49.7 nm for the four populations, *n* = 79, 114, 134, and 95 particles, respectively. Scale bars: 250 nm for PS and 500 nm for PMMA.

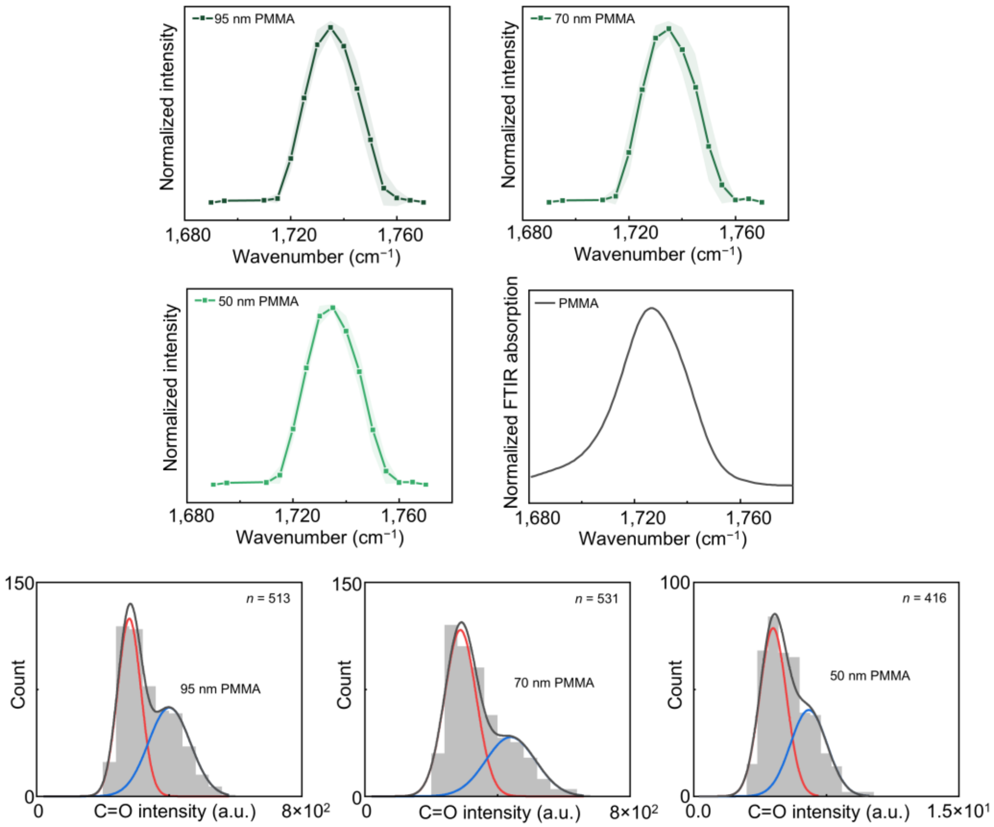


**Supplementary Fig. 10 Averaged IR-AMES spectra and C=O intensity distributions of 95-nm, 70-nm and 50-nm PMMA nanoparticles.** Top: Averaged IR-AMES spectra of PMMA particles, together with the bulk FTIR absorption spectrum of PMMA. Solid line: mean spectra, shaded region: standard deviation. Bottom: Histograms of the C=O band intensity at 1,735 $cm^{-1}$, fitted with Gaussian components. The red Gaussian corresponds to single particles, while the blue Gaussian represents multi-particle aggregates or multiple particles located within sub-diffraction distances.

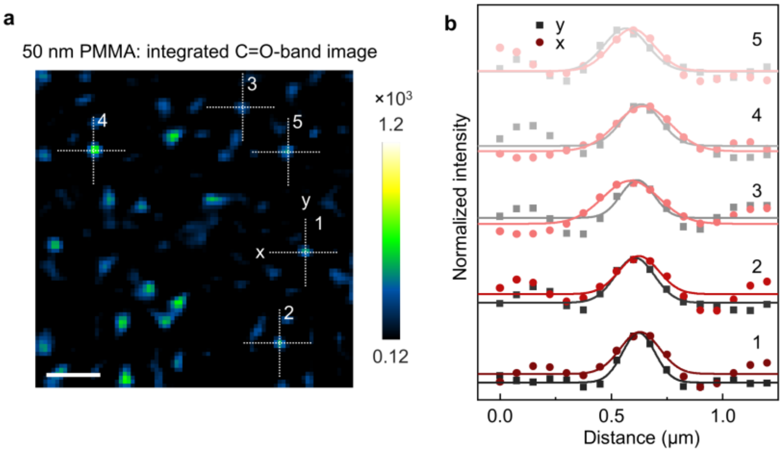


**Supplementary Fig. 11 Representative FWHM analysis of 50-nm PMMA particles. a,** IR-AMES image of 50-nm PMMA particles integrated over the C=O band (1,715–1,755 $cm^{-1}$). Scale bar, 1 μm. **b,** Corresponding normalized intensity profiles along the x and y directions. Solid lines show Gaussian fits used to determine the FWHM values. Dashed lines in **a** indicate the *x*- and *y*-direction line profiles through five representative particles used for FWHM analysis.

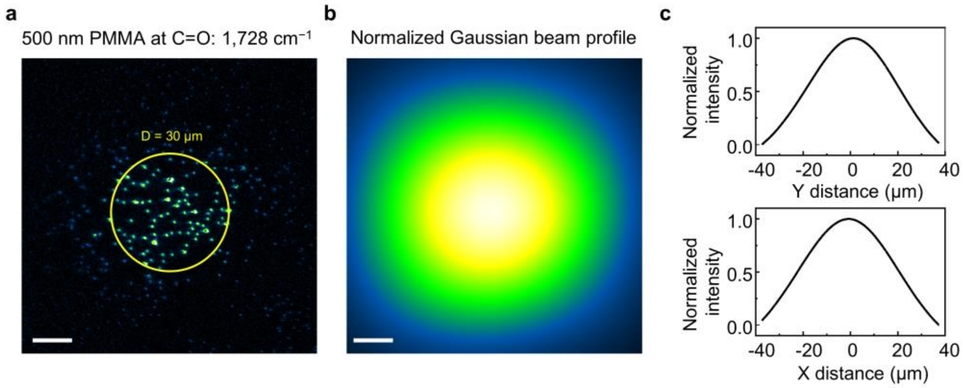


**Supplementary Fig. 12 IR beam-shape fitting and image-area selection. a,** IR-AMES image of 500-nm PMMA at the C=O band (1,728 $cm^{-1}$). **b,** The IR illumination profile is fitted with a 2D Gaussian function. **c,** Line profiles show a beam FWHM of 50 µm (*y*-axis) and 45.7 µm (*x*-axis), corresponding to an elliptical illumination area with semi-major axis $a$ = 25 µm and semi-minor axis $b$ = 22.85 µm. A uniform central region with diameter $D$ = 30 µm is selected as the effective imaging area for all measurements. Scale bars: 10 µm.

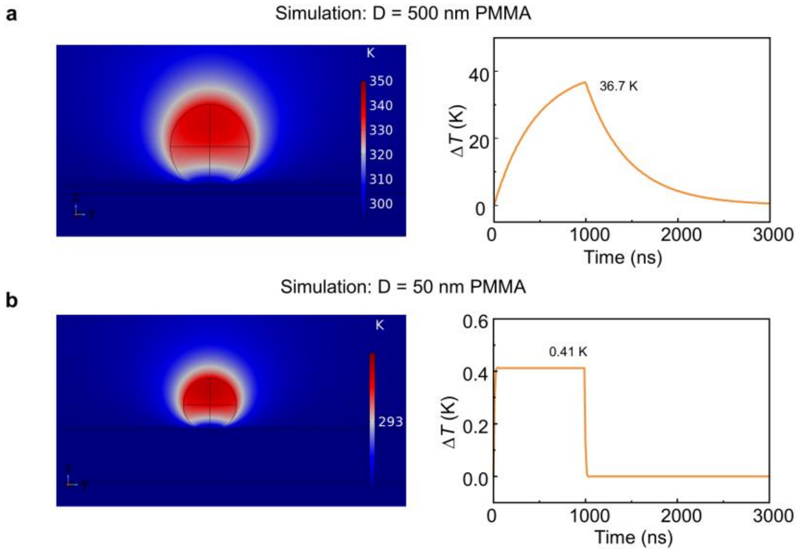


**Supplementary Fig. 13 Simulated temperature distribution and transient heating response in the IR-AMES system. a,** Simulated temperature distribution and transient temperature rise of a 500-nm PMMA bead in air under a single 1-µs IR pulse. The particle reaches a peak temperature increase of ~36.7 K before cooling. **b,** Corresponding simulation for a 50-nm PMMA bead, showing a smaller temperature rise of ~0.41 K under the same IR pulse.

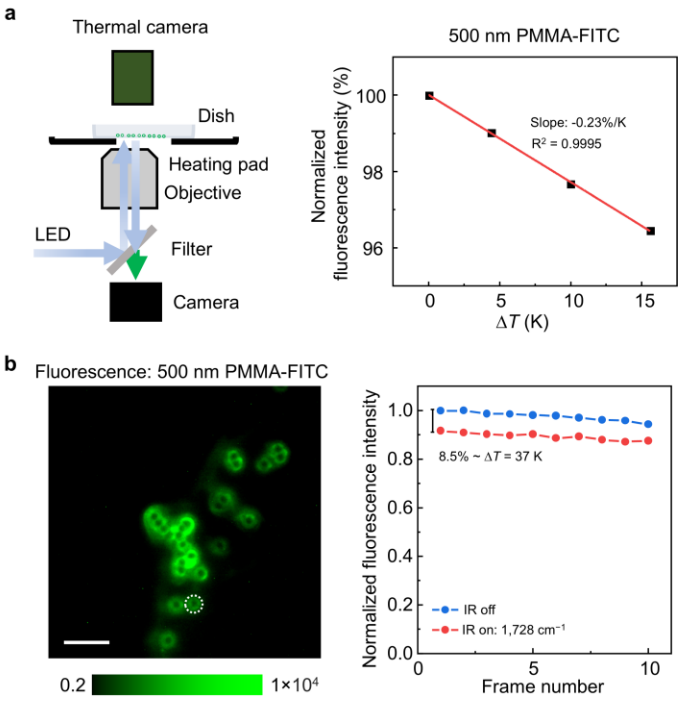


**Supplementary Fig. 14 Experimental temperature response of 500-nm PMMA particles. a,** Schematic of the widefield fluorescence setup used to measure temperature-dependent fluorescence of 500-nm PMMA-FITC beads in air, together with the calibration curve of normalized fluorescence intensity versus temperature change ($\Delta T$). Fluorescence decreases linearly with temperature increase, with a slope of −0.23% per K. **b,** Fluorescence image of 500-nm PMMA-FITC beads and the corresponding normalized fluorescence intensity modulation under IR-off and IR-on (1,728 $cm^{-1}$) conditions. An 8.5% fluorescence decrease corresponds to a temperature rise of ~37 K on single particle. Scale bar: 5 µm.

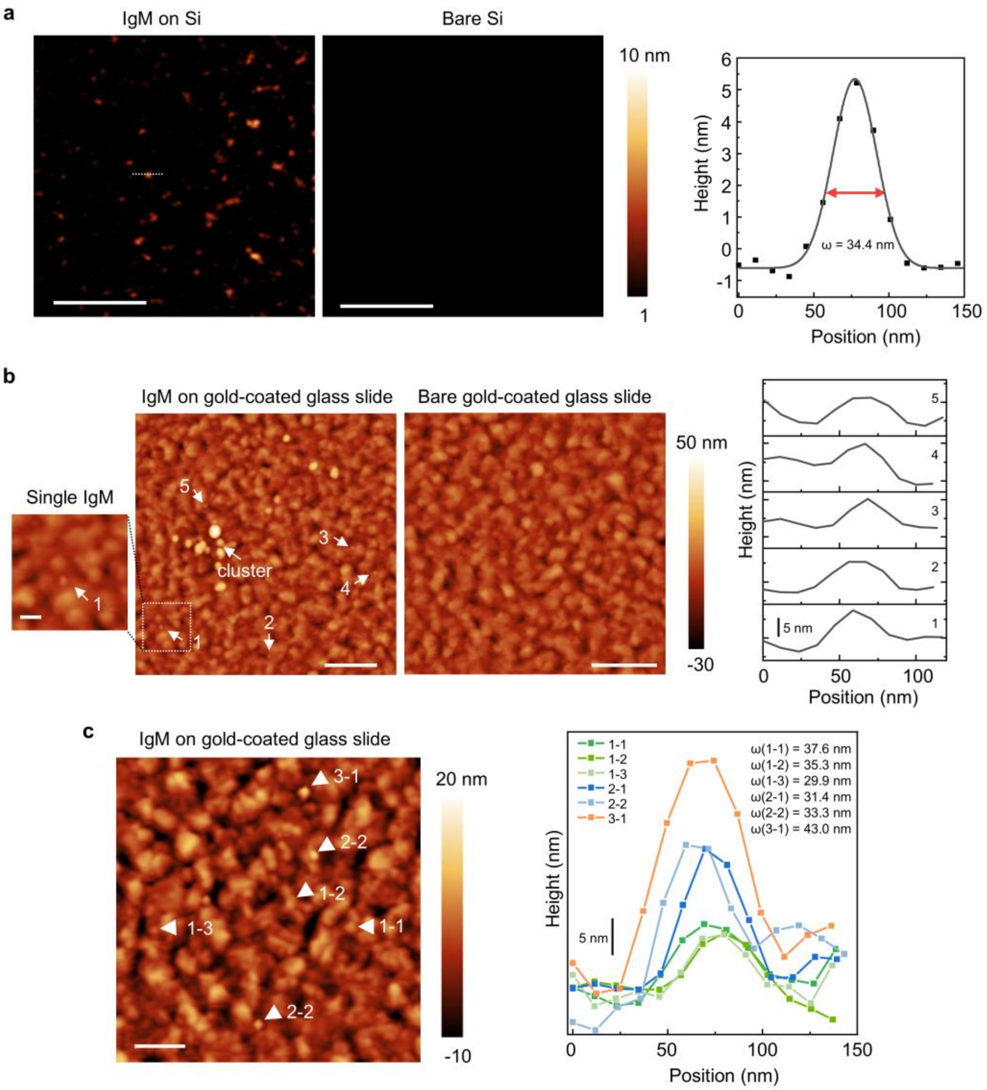


**Supplementary Fig. 15 AFM characterization of individual IgM complexes and small aggregates. a,** AFM images of IgM deposited on a flat Si substrate and a bare Si control. Individual IgM complexes are resolved as isolated particles on Si, whereas the bare substrate remains featureless. The representative line profile shows a particle height of approximately 5 nm and a Gaussian-fitted lateral FWHM of 34.4 nm. Scale bars: 250 nm. **b,** AFM images of IgM deposited on a gold-coated glass slide and of a bare gold-coated slide. The bare gold surface exhibits nanoscale roughness, whereas additional isolated particle-like features and clustered regions are observed after IgM deposition. Representative height profiles of particles 1–5 are shown on the right. Scale bars: 500 nm; inset scale bar: 100 nm. **c,** AFM characterization of IgM on a gold-coated glass slide prepared from the same sample used for IR-AMES measurements in **Fig. 2a**. The AFM image was acquired from a separate field of view and is not spatially co-localized with the IR-AMES image. Representative height profiles distinguish single IgM complexes (1-1 to 1-3) from double (2-1 and 2-2) and triple (3-1) IgM aggregates. Scale bar: 250 nm.

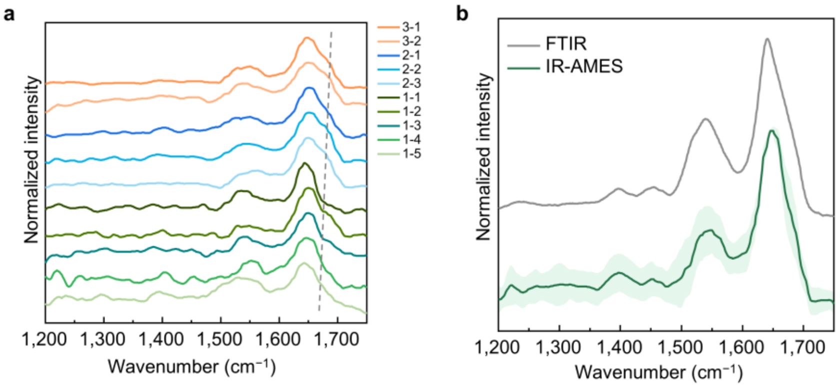


**Supplementary Fig. 16 Validation of IgM spectral fidelity over the fingerprint region. a,** IR-AMES spectra of individual IgM complexes and small aggregates identified in **Fig. 2a** over the 1,200–1,750 $cm^{-1}$ spectral range. Spectra were smoothed for display. Grey dashed lines indicate spectral regions where differences between individual IgM complexes and small aggregates are apparent. **b,** Comparison of the mean IR-AMES spectrum with the bulk FTIR spectrum of dried IgM. Solid lines: mean spectra, shaded regions: standard deviation.

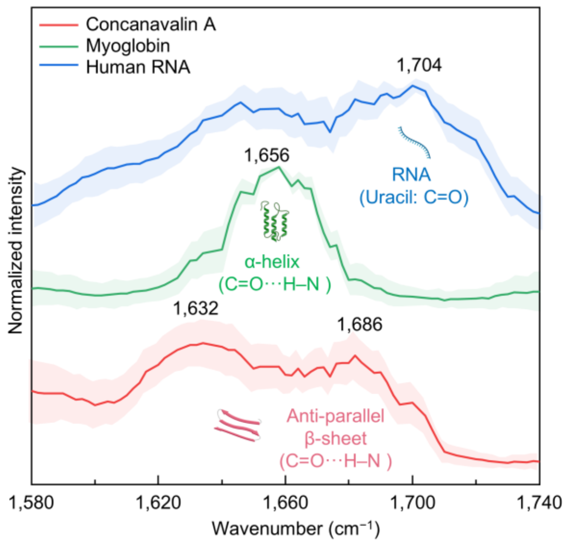


**Supplementary Fig. 17 IR-AMES spectra of reference biomolecules.** Averaged IR-AMES spectra of concanavalin A, myoglobin and human RNA in solution. Characteristic spectral features are observed near 1,632 and 1,686 $cm^{-1}$ for anti-parallel β-sheet-rich concanavalin A, near 1,656 $cm^{-1}$ for α-helix-rich myoglobin and near 1,704 $cm^{-1}$ for the uracil C=O vibration of RNA. Solid lines: mean spectra, shaded regions: standard deviation ($n$ = 40 particles per group). Spectra were normalized to 0–1 and vertically offset for display.

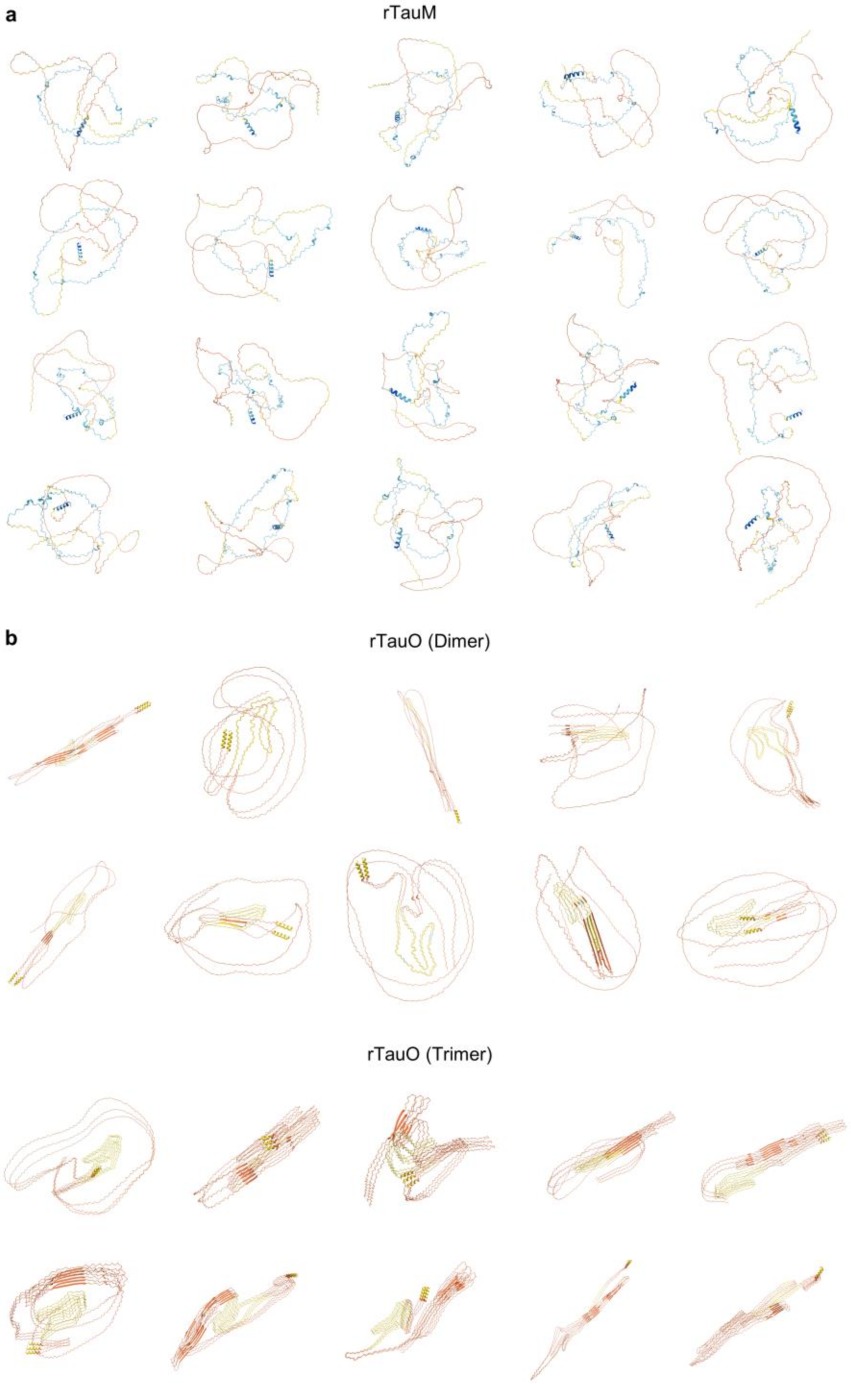


**Supplementary Fig. 18 AlphaFold3 structure predictions of recombinant 2N4R tau monomers (rTauM) and oligomers (rTauO). a,** rTauM are structurally heterogeneous due to their intrinsically disordered nature, showing random-coil-dominant conformations and short α-helix structures. **b,** Many dimers or trimers rapidly adopt β-sheet-rich structures within disordered conformations, leading to coexistence of β-sheet–rich, partially folded, and highly disordered conformational states within early tau oligomers. Given the intrinsically disordered nature of tau, the prediction confidence is variable, and these models are regarded as structural hypotheses.

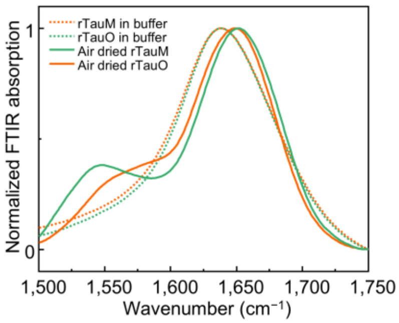


**Supplementary Fig. 19 FTIR spectra of recombinant tau monomer (rTauM) and oligomer (rTauO) samples in buffer and after drying.** In solution, strong water absorption obscures the protein amide-I features. Drying recovers the protein vibrational bands, whereas ensemble averaging limits the ability to resolve structural heterogeneity within the rTauM and rTauO populations.

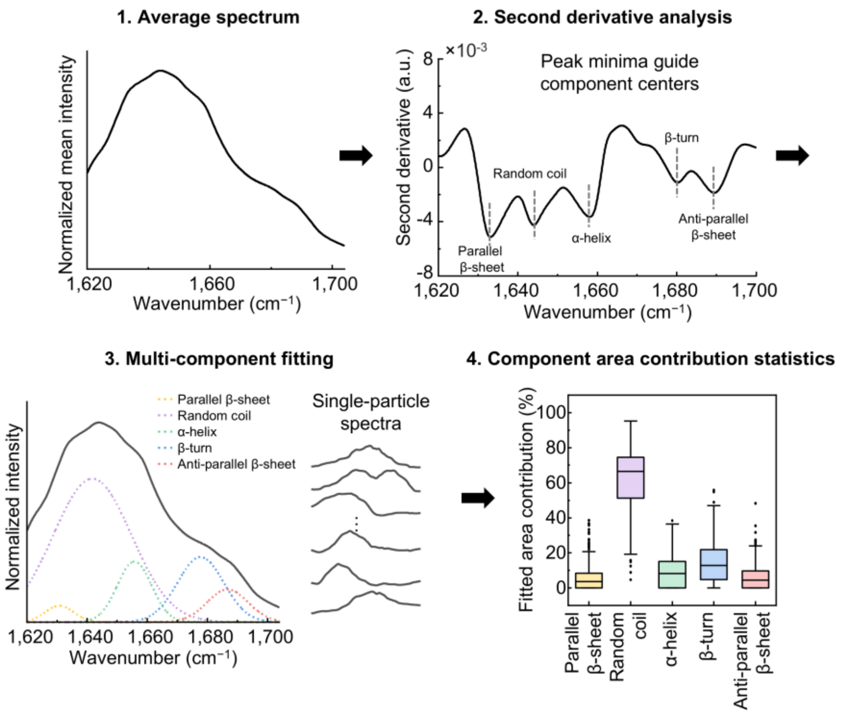


**Supplementary Fig. 20 Workflow for component analysis of single-particle IR-AMES spectra. 1,** The raw single-particle spectra are averaged within each sample group, and the resulting mean spectrum is smoothed using a Savitzky–Golay filter. **2,** Second-derivative analysis is used to identify characteristic spectral minima and guide the assignment of component centers. **3,** Individual single-particle spectra are fitted using a constrained multi-component model, with component positions guided by the second-derivative analysis. **4,** The integrated area of each fitted component is calculated for every particle and used to determine the distributions of component contributions across the population.

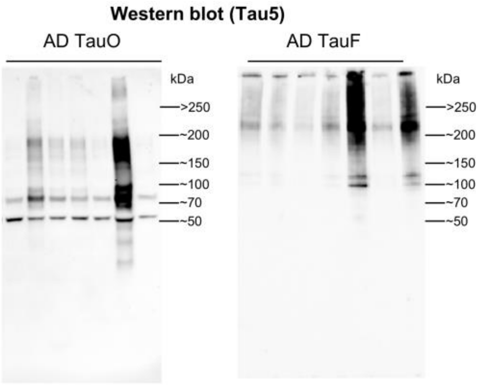


**Supplementary Fig. 21 Western blot analysis of human brain-derived AD TauO and TauF.** Native, non-reducing and detergent-free Western blot analysis using the Tau5 antibody. Each lane represents an individual patient sample ($n$ = 7 per group). AD TauO shows predominantly lower-molecular-weight Tau-positive species, whereas AD TauF contains a greater proportion of high-molecular-weight species (> 200 kDa). The AD TauO and AD TauF samples used in **Fig. 4a** were prepared by pooling five patient samples selected based on the amount of tau recovered during extraction.

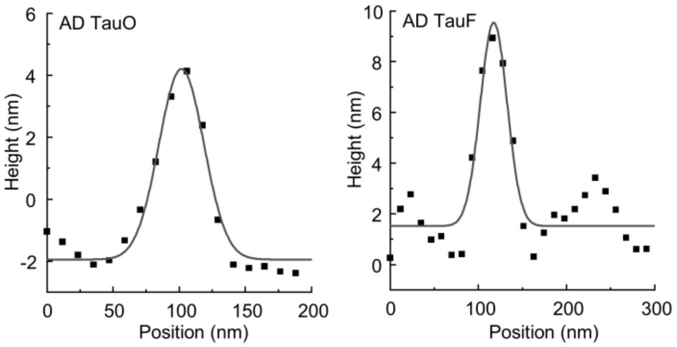


**Supplementary Fig. 22 Size measurements of AD TauO and AD TauF.** Height profiles with Gaussian fittings (solid lines), data extracted from AFM images in **Fig. 4a**.

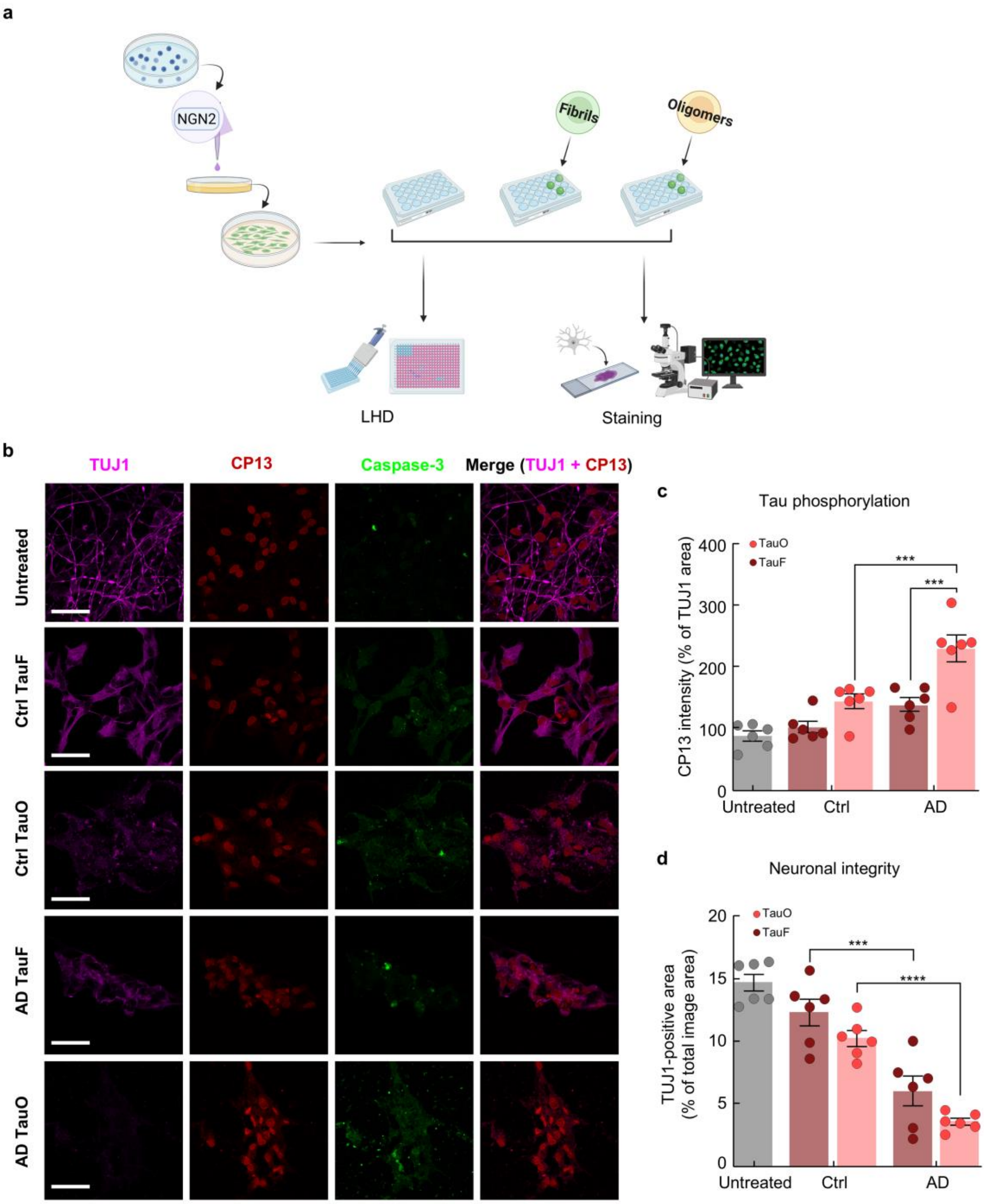


**Supplementary Fig. 23 Effects of human tau species on iPSC-derived neurons. a,** Schematic of the experimental design. **b,** Representative fluorescence images of iPSC-derived neurons untreated and treated with different human tau species for 24 h. Scale bars: 50 µm. **c-d,** Quantification of tau phosphorylation by CP13 intensity (**c**) and neuronal integrity by TUJ1 intensity (**d**). $n$ = 6. Data were expressed as mean ± s.d. Column means were compared using two-way ANOVA, with ***p < 0.001 and ****p < 0.0001.

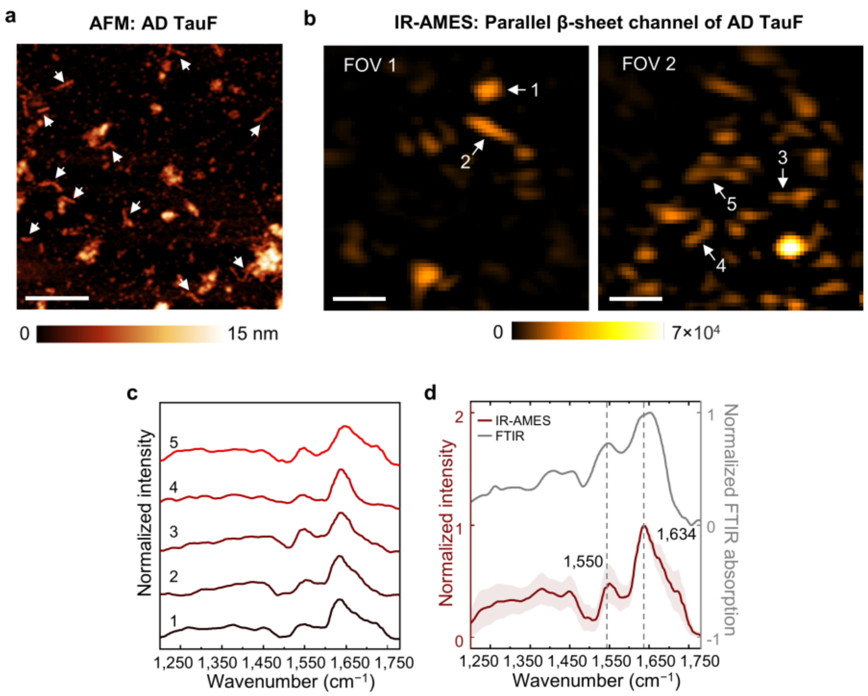


**Supplementary Fig. 24 Morphological and spectroscopic characterization of human brain-derived tau fibrils. a,** Representative AFM image of AD TauF, showing elongated fibrillar structures (arrows). **b,** Representative IR-AMES images of AD TauF acquired in the parallel β-sheet channel from two fields of view (FOV). **c,** Representative IR-AMES spectra extracted from the structures indicated in **b**. **d,** Comparison of the mean IR-AMES spectrum of AD TauF with the ensemble FTIR spectrum. Solid lines: mean spectra, shaded regions: standard deviation. Both spectra show a prominent parallel β-sheet-associated band near 1,634 $cm^{-1}$. Scale bars: 1 μm.

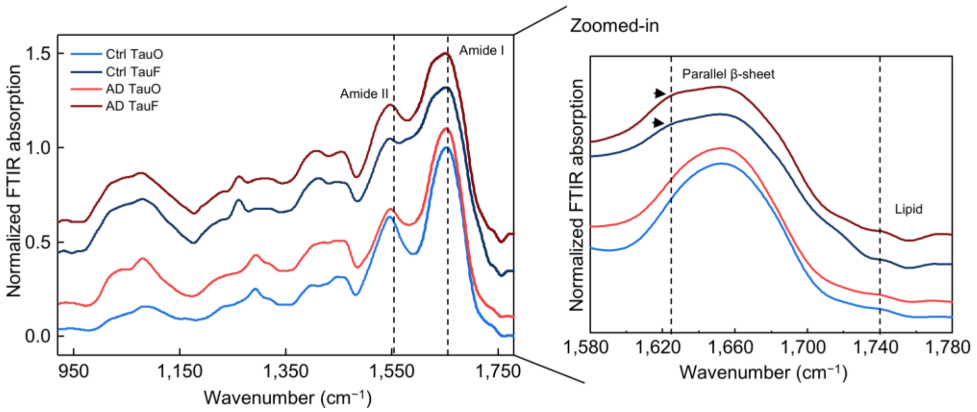


**Supplementary Fig. 25 FTIR spectra of dried human-derived tau samples.** Full fingerprint-region FTIR spectra of Ctrl TauO, AD TauO, Ctrl TauF and AD TauF (left), with an expanded view of the amide-I region (right). All samples show characteristic protein absorption bands, including amide II (~1,550 $cm^{-1}$) and amide I (~1,650 $cm^{-1}$). Both AD TauO and Ctrl TauO exhibit strong amide-I absorption, with no obvious spectral differences between the two oligomer populations under dried conditions. In contrast, AD TauF and Ctrl TauF show a distinct peak consistent with parallel β-sheet structures. All tau samples show only weak lipid-associated signals, likely reflecting partial loss or removal of lipid species during extraction and purification steps.

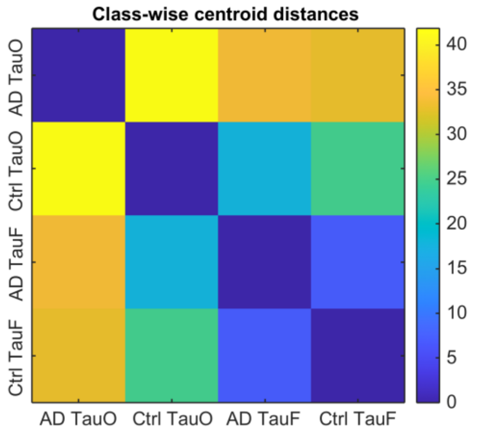


**Supplementary Fig. 26 Centroid distances in t-SNE space.** Euclidean distances between class centroids calculated from the embedding shown in **Fig. 4l.** AD TauF and Ctrl TauF exhibit the smallest inter-class distance, whereas AD TauO is most separated from other tau species. Values represent relative proximity in embedding space (arbitrary units).

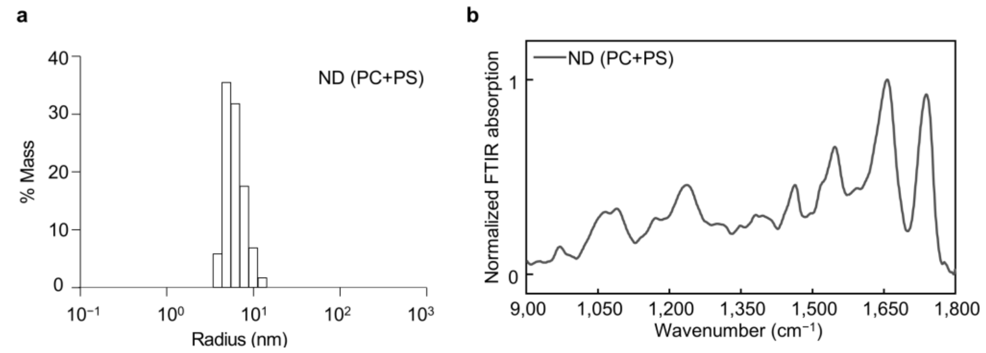


**Supplementary Fig. 27 Characterization of nanodiscs. a,** Radius distribution of ND (PC/PS) measured by dynamic light scattering. **b,** FTIR spectrum of bulk ND (PC/PS), showing characteristic protein- and lipid-associated vibrational bands consistent with their major components.

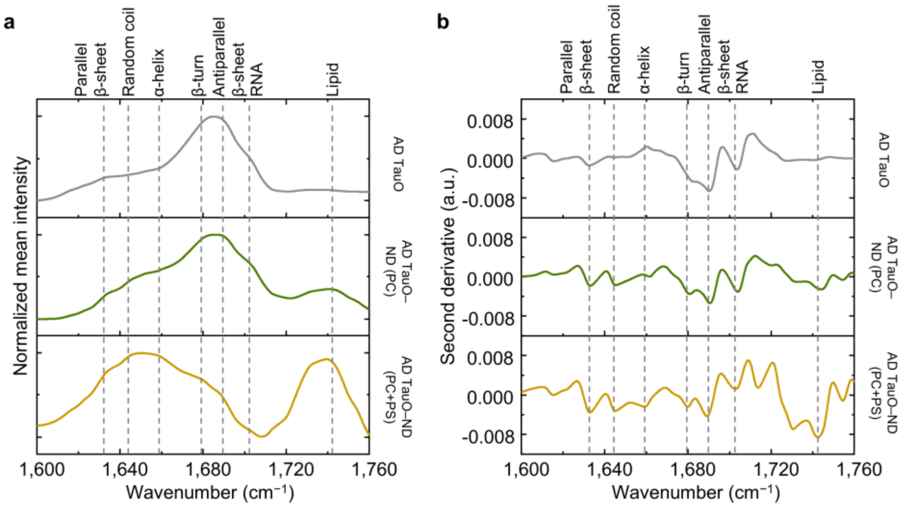


**Supplementary Fig. 28 Amide-I spectral features and second-derivative analysis of AD TauO-associated nanodisc samples. a,** Mean amide-I spectra of AD TauO, AD TauO-ND (PC) and AD TauO-ND (PC+PS). **b,** Corresponding second-derivative spectra, highlighting spectral minima associated with parallel β-sheet, random coil, α-helix, β-turn, antiparallel β-sheet, RNA- and lipid-related vibrational features. Dashed lines indicate reference spectral positions used to guide component-center assignment for subsequent spectral decomposition.

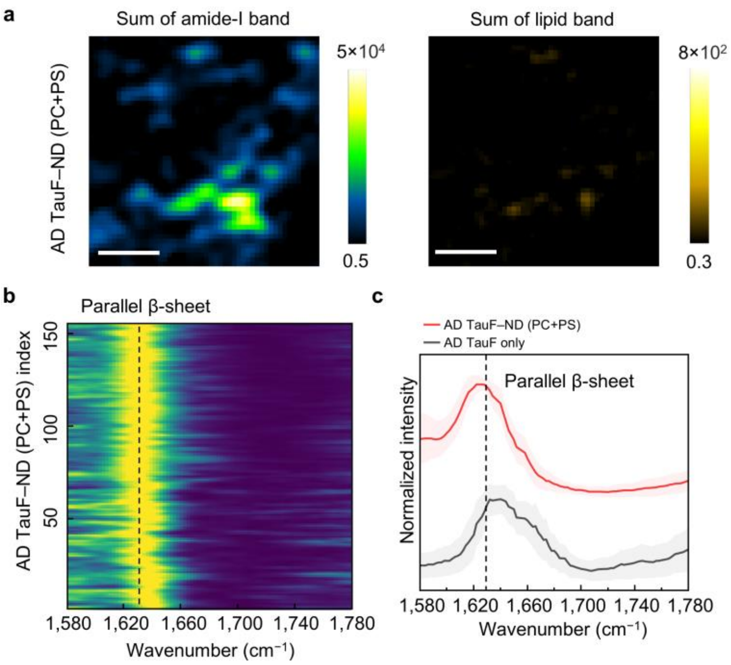


**Supplementary Fig. 29 IR-AMES images of AD TauF–ND complex. a,** IR-AMES images of AD TauF following ND (PC+PS) incubation. Top row: integrated amide-I signal; bottom row: integrated lipid signal. Scale bars, 1 μm. **b,** Heatmaps of IR-AMES spectra from AD TauF–ND (PC+PS) complexes. Spectra were normalized to 0–1. **c,** Average spectra of AD TauF–ND (PC+PS) complexes in **b** (red, $n$ = 155) and average spectra of AD TauF only (gray, $n$ = 150). Solid lines: mean spectra, shaded regions: standard deviation. Spectra were normalized to 0–1 and vertically offset for display.

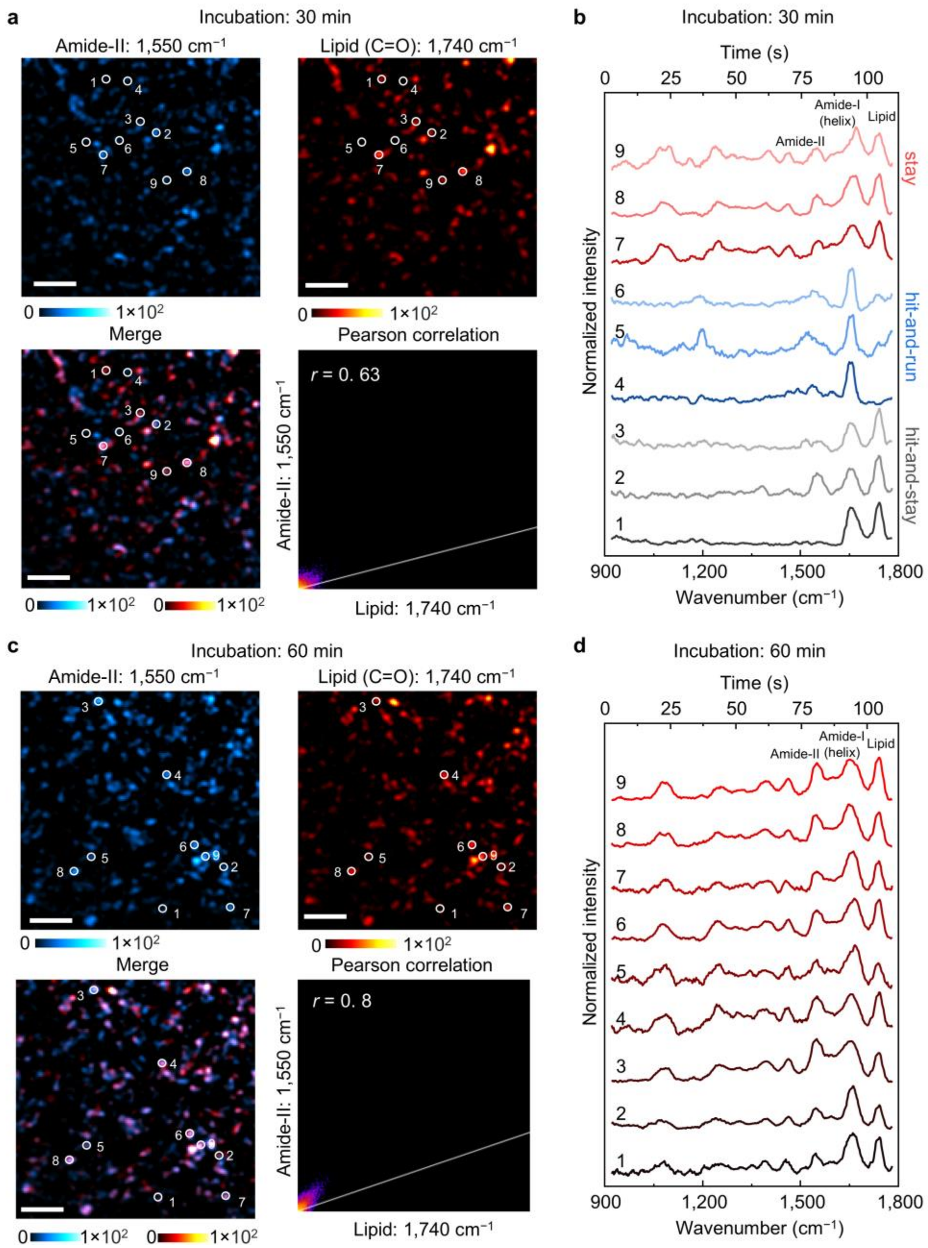


**Supplementary Fig. 30 Control experiments excluding motion-induced artifacts in solution.** **a**, **b**, IR-AMES measurements of ND (PC+PS) (150 nM) after 30 min incubation. **a**, IR-AMES images at the amide-II (1,550 $cm^{-1}$) and lipid (1,740 $cm^{-1}$) channels, corresponding merged image, and Pearson correlation map ($r = 0.63$). **b**, IR-AMES spectra from representative nanoparticles marked in **a**, showing heterogeneous measurement of transient and partially stabilized landing events. **c**, **d**, IR-AMES measurements of ND (PC+PS) (150 nM) after 60 min incubation. **c**, IR-AMES images at the amide-II and lipid channels, merged image, and Pearson correlation map ($r = 0.80$), demonstrating strong spatial co-localization between protein and lipid signals. **d**, IR-AMES spectra from representative nanoparticles marked in **c**, exhibiting stable and reproducible spectral features throughout acquisition, consistent with surface-confined nanodiscs and excluding motion-induced artifacts.

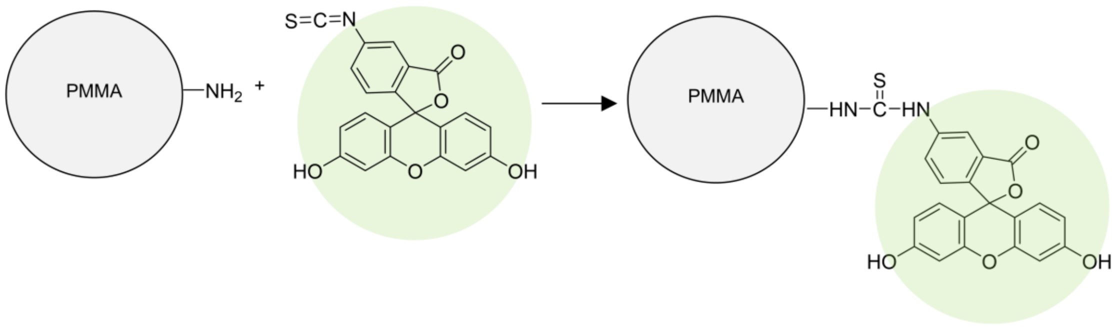


**Supplementary Fig. 31 Synthetic procedure of PMMA-FITC beads.** Amine-functionalized 500-nm PMMA beads were covalently labeled with FITC via isothiocyanate group, where the fluorescein core provides temperature-sensitive green fluorescence.

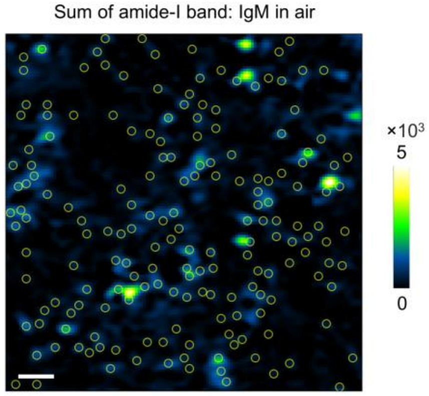


**Supplementary Fig. 32 Single-particle region-of-interest (ROI) selection for IR-AMES spectral analysis.** Image of single IgM in air is used as the example. An integrated IR-AMES image of IgM molecules was generated by summing the hyperspectral images over the amide-I band (1,620–1,660 $cm^{-1}$). Individual particles were automatically detected using TrackMate in ImageJ. Yellow circles indicate diffraction-limited ROIs (3-pixel diameter, corresponding to ~225 nm). Mean intensities within each ROI were extracted at each wavenumber to construct single-particle IR-AMES spectra. Scale bar: 1 µm.

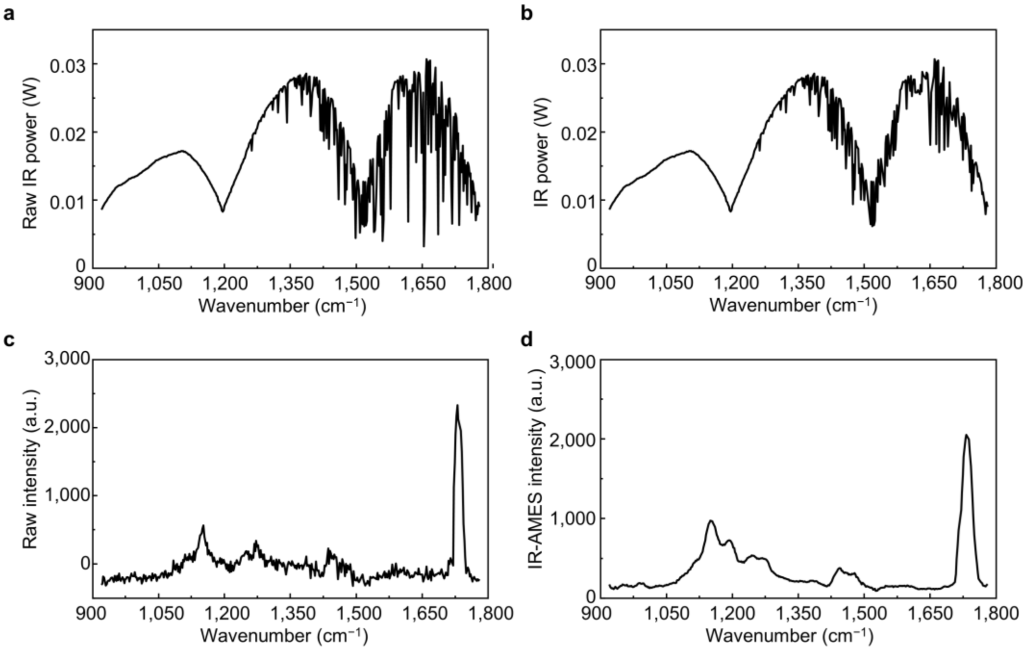


**Supplementary Fig. 33 IR-AMES spectral processing. a,** Raw average IR power measured at the sample plane. Sharp negative spikes in the curve arise from water vapor absorption. **b,** Average IR power spectrum after excluding wavenumbers strongly affected by water vapor absorption, used for spectral normalization. **c**, Raw IR-AMES spectrum extracted from a single 500-nm PMMA particle. **d**, IR-AMES spectrum of the same particle after spectral normalization.

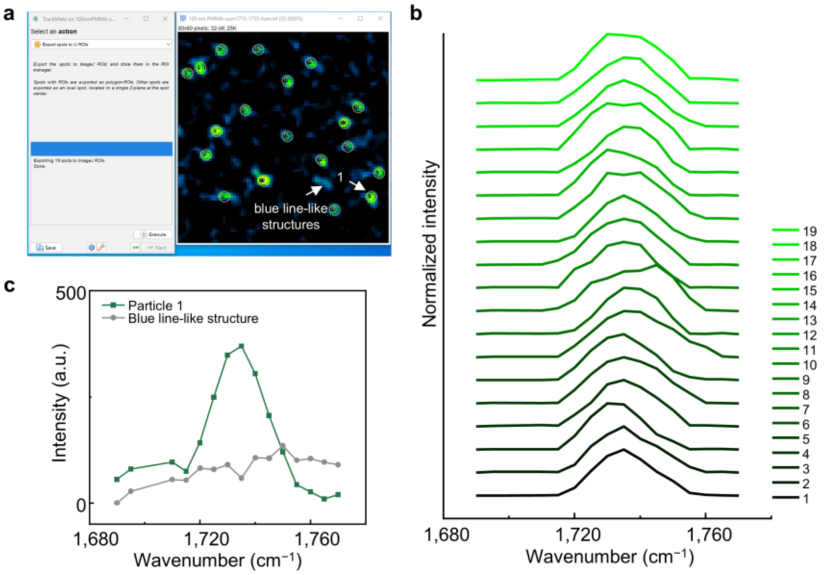


**Supplementary Fig. 34 Spectroscopic validation of automated nanoparticle identification. a,** Automated identification of diffraction-limited spots in an integrated C=O-band IR-AMES image of 95-nm PMMA nanoparticles in ImageJ. **b,** IR-AMES spectra extracted from automatically identified particles, showing the characteristic PMMA C=O resonance near 1,730 $cm^{-1}$. **c,** Representative spectra from an identified PMMA particle and a blue line-like background feature. The particle shows the expected C=O peak, whereas the background feature lacks the PMMA-specific vibrational signature.
